# A neurotrophin functioning with a Toll regulates structural plasticity in a dopaminergic circuit

**DOI:** 10.1101/2023.01.04.522695

**Authors:** Jun Sun, Francisca Rojo-Cortés, Suzana Ulian-Benitez, Manuel G. Forero, Guiyi Li, Deepanshu Singh, Xiaocui Wang, Sebastian Cachero, Marta Moreira, Dean Kavanagh, Gregory Jefferis, Vincent Croset, Alicia Hidalgo

**Author notes:** Author for correspondence: 00 44 (0)121 4145416.

## Abstract

Experience shapes the brain, as neural circuits can be modified by neural stimulation or the lack of it. The molecular mechanisms underlying structural circuit plasticity and how plasticity modifies behaviour, are poorly understood. Subjective experience requires dopamine, a neuromodulator that assigns a value to stimuli, and it also controls behaviour, including locomotion, learning and memory. In *Drosophila*, Toll receptors are ideally placed to translate experience into structural brain change. *Toll-6* is expressed in dopaminergic neurons (DANs), raising the intriguing possibility that Toll-6 could regulate structural plasticity in dopaminergic circuits. *Drosophila* neurotrophin-2 (DNT-2) is the ligand for Toll-6 and Kek-6, but whether it is required for circuit structural plasticity was unknown. Here, we show that *DNT-2* expressing neurons connect with DANs, and they modulate each other. Loss of function for *DNT-2* or its receptors *Toll-6* and kinase-less Trk-like *kek-6* caused DAN and synapse loss, impaired dendrite growth and connectivity, decreased synaptic sites and caused locomotion deficits. By contrast, over-expressed *DNT-2* increased DAN cell number, dendrite complexity and promoted synaptogenesis. Neuronal activity modified DNT-2, it increased synaptogenesis in DNT-2-positive neurons and DANs, and over-expression of DNT-2 did too. Altering the levels of DNT-2 or Toll-6 also modified dopamine-dependent behaviours, including locomotion and long-term memory. To conclude, a feedback loop involving dopamine and DNT-2 sculpted the circuits engaged, and DNT-2 with Toll-6 and Kek-6 induced structural plasticity in this circuit modifying brain function and behaviour.

## INTRODUCTION

The brain can change throughout life, as new cells are formed or eliminated, axonal and dendritic arbours can grow or shrink, synapses can form or be eliminated (Wiesel 1982, Feldman and Brecht 2005, Holtmaat and Svoboda 2009, Gage 2019). Such changes can be driven by experience, that is, neuronal activity or the lack of it (Wiesel 1982, Maguire et al. 2000, Cotman and Berchtold 2002, Feldman and Brecht 2005, Sur and Rubenstein 2005, Holtmaat and Svoboda 2009, Woollett and Maguire 2011, Chen and Brumberg 2021, Bharmauria et al. 2022). Structural changes result in remodelling of connectivity patterns, and these bring about modifications of behaviour. These can be adaptive, dysfunctional or simply the consequence of opportunistic connections between neurons (Kuner and Flor 2016, Leemhuis et al. 2019, Yang et al. 2020). It is critical to understand how structural modifications to cells influence brain function. This requires linking with cellular resolution molecular mechanisms, neural circuits and resulting behaviours.

In the mammalian brain, the neurotrophins (NTs: BDNF, NGF, NT3, NT4) are growth factors underlying structural brain plasticity (Poo 2001, Lu et al. 2005, Park and Poo 2013). They promote neuronal survival, connectivity, neurite growth, synaptogenesis, synaptic plasticity and Long-Term Potentiation (LTP), via their Trk and p75^NTR^ receptors (Poo 2001, Lu et al. 2005, Park and Poo 2013). In fact, all anti-depressants function by stimulating production of BDNF and signalling via its receptor TrkB, leading to increased brain plasticity (Casarotto et al. 2021, Castren and Monteggia 2021). Importantly, NTs have dual functions and can also induce neuronal apoptosis, neurite loss, synapse retraction and Long-Term Depression (LTD), via p75^NTR^ and Sortilin (Lu et al. 2005). Remarkably, these latter functions are shared with neuroinflammation, which in mammals involves Toll-Like Receptors (TLRs)(Squillace and Salvemini 2022). TLRs and Tolls have universal functions in innate immunity across the animals (Gay and Gangloff 2007), and consistently with this, TLRs in the CNS are mostly studied in microglia. However, mammalian *TLRs* are expressed in all CNS cell types, where they can promote not only neuroinflammation, but also neurogenesis, neurite growth and synaptogenesis and regulate memory – independently of pathogens, cellular damage or disease (Ma et al. 2006, Rolls et al. 2007, Okun et al. 2010, Okun et al. 2011, Patel et al. 2016, Chen, CY et al. 2019). Whether TLRs have functions in structural brain plasticity and behaviour remains little explored, and whether they can function together with neurotrophins in the mammalian brain is unknown.

Progress linking cellular and molecular events to circuit and behavioural modification has been rather daunting and limited using mammals (Wang et al. 2022). The *Drosophila* adult brain is plastic and can be modified by experience and neuronal activity (Technau 1984, Barth and Heisenberg 1997, Barth et al. 1997, Sachse et al. 2007, Kremer et al. 2010, Sugie et al. 2015, Linneweber et al. 2020, Baltruschat et al. 2021, Coban et al. 2024). Different living conditions, stimulation with odorants or light, circadian rhythms, nutrition, long-term memory and experimentally activating or silencing neurons, modify brain volume, alter circuit and neuronal shape, and remodel synapses, revealing experience-dependent structural plasticity (Heisenberg et al. 1995, Barth and Heisenberg 1997, Barth et al. 1997, Devaud et al. 2001, Gorska-Andrzejak et al. 2005, Sachse et al. 2007, Fernandez et al. 2008, Kremer et al. 2010, Bushey et al. 2011, Sugie et al. 2015, Duhart et al. 2020, Baltruschat et al. 2021, Vaughen et al. 2022, Coban et al. 2024). Furthermore, the *Drosophila* brain is also susceptible to neurodegeneration (Bolus et al. 2020). However, the molecular and circuit mechanisms underlying structural brain plasticity are mostly unknown in *Drosophila*.

*Toll* receptors are expressed across the *Drosophila* brain, in distinct but overlapping patterns that mark the anatomical brain domains (Li et al. 2020). Tolls share a common signalling pathway downstream, that can drive at least four distinct cellular outcomes – cell death, survival, quiescence, proliferation – depending on context (McIlroy et al. 2013, Foldi et al. 2017, Anthoney et al. 2018, Li et al. 2020). They are also required for connectivity and structural synaptic plasticity and they can also induce cellular events independently of signalling (McIlroy et al. 2013, Ward et al. 2015, McLaughlin et al. 2016, Foldi et al. 2017, Ulian-Benitez et al. 2017, Li et al. 2020). These nervous system functions occur in the absence of tissue damage or infection. This is consistent with the fact that - as well as universal functions in innate immunity - Tolls also have multiple non-immune functions also outside the CNS, including the original discovery of Toll in dorso-ventral patterning, cell intercalation, cell competition and others (Meyer et al. 2014, Pare et al. 2014, Anthoney et al. 2018, Tamada et al. 2021). The Toll distribution patterns in the adult brain and their ability to switch between distinct cellular outcomes means they are ideally placed to translate experience into structural brain change (Li et al. 2020).

We had previously observed that in the adult brain, *Toll-6* is expressed in dopaminergic neurons (McIlroy et al. 2013). Dopamine is a key neuromodulator that regulates wakefulness and motivation, experience valence, such as reward, and it is essential for locomotion, learning and memory (Riemensperger et al. 2011, Waddell 2013, Adel and Griffith 2021). In *Drosophila*, Dopaminergic neurons (DANs) form an associative neural circuit together with mushroom body Kenyon cells (KCs), Dorsal Anterior Lateral neurons (DAL) and Mushroom Body Output neurons (MBONs) (Chen et al. 2012, Aso et al. 2014a, Boto et al. 2020, Adel and Griffith 2021). Kenyon cells receive input from projection neurons of the sensory systems, they then project through the mushroom body lobes where they are intersected by DANs to regulate MBONs to drive behaviour (Heisenberg 2003, Aso et al. 2014b, Boto et al. 2020). This associative circuit is required for learning, long-term memory and goal-oriented behaviour (Chen et al. 2012, Guven-Ozkan and Davis 2014, Adel and Griffith 2021). During experience, involving sensory stimulation from the external world and from own actions, dopamine assigns a value to otherwise neutral stimuli, labelling the neural circuits engaged (Boto et al. 2020). Thus, this raises the possibility that a link of Toll-6 to dopamine could enable translating experience into circuit modification to modulate behaviour.

In *Drosophila*, Toll receptors can function both independently of ligand-binding and by binding Spätzle (Spz) protein family ligands, also known as *Drosophila* neurotrophins (DNTs), that are sequence, structural and functional homologues of the mammalian NTs (DeLotto and DeLotto 1998, Weber et al. 2003, Hoffmann et al. 2008a, Hoffmann et al. 2008b, Zhu et al. 2008, Lewis et al. 2013, McIlroy et al. 2013, Foldi et al. 2017). Like mammalian NTs, DNTs also promote cell survival, connectivity, synaptogenesis and structural synaptic plasticity, and they can also promote cell death, depending on context (Zhu et al. 2008, McIlroy et al. 2013, Sutcliffe et al. 2013, Foldi et al. 2017, Ulian-Benitez et al. 2017). As well as Tolls, DNTs are also ligands for Kekkon (Kek) receptors, kinase-less homologues of the mammalian NT Trk receptors and are required for structural synaptic plasticity (Ulian-Benitez et al. 2017). Importantly, the targets regulated by Tolls and Keks - ERK, NFκB, PI3K, JNK, CaMKII – are shared with those of mammalian NT receptors Trk and p75^NTR^, and have key roles in structural and functional plasticity across the animals (Park and Poo 2013, Foldi et al. 2017, Ulian-Benitez et al. 2017, Yang et al. 2020, Tamada et al. 2021).

Here, we focus on Drosophila neurotrophin −2 (DNT-2), proved to be the ligand of Toll-6 and Kek-6, with in vitro, cell culture and in vivo evidence (McIlroy et al. 2013, Foldi et al. 2017, Ulian-Benitez et al. 2017). Here we asked how DNT-2, Toll-6 and Kek-6 are functionally related to dopamine, whether they and neuronal activity – as a proxy for experience - can modify neural circuits, and how structural circuit plasticity modifies dopamine-dependent behaviours.

## RESULTS

### DNT-2A, Toll-6 and Kek-6 neurons are integrated in a dopaminergic circuit

To allow morphological and functional analyses of *DNT-2* expressing neurons, we generated a *DNT-2Gal4* line using CRISPR/Cas9 and drove expression of the membrane-tethered-GFP *FlyBow1.1* reporter. We identified at least 12 DNT-2+ neurons and we focused on four anterior DNT-2A neurons per hemi-brain (Figure 1A, B). Using the post-synaptic marker Denmark, DNT-2A dendrites were found at the prow (PRW) and flange (FLA) region, whereas axonal terminals visualised with the pre-synaptic marker synapse defective 1 (Dsyd1-GFP) resided at the superior medial protocerebrum (SMP) (Figure 1C, C’’). We additionally found post-synaptic signal at the SMP and pre-synaptic signal at the FLA/PRW (Figure 1C, C’), suggesting bidirectional communication at both sites. Using Multi-Colour Flip-Out (MCFO) to label individual cells stochastically (Nern et al. 2015, Costa et al. 2016), single neuron clones revealed variability in the DNT-2A projections across individual flies (Figure 1D), consistently with developmental and activity-dependent structural plasticity in *Drosophila* (Heisenberg et al. 1995, Kremer et al. 2010, Sugie et al. 2015, Mayseless et al. 2018, Li et al. 2020, Linneweber et al. 2020, Baltruschat et al. 2021). We found that DNT-2A neurons are glutamatergic as they express the vesicular glutamate transporter vGlut (Figure 1E, Figure S1A) and lack markers for other neurotransmitter types (Figure S1). DNT-2A terminals overlapped with those of dopaminergic neurons (Figure 1G), suggesting they could receive inputs from neuromodulatory neurons. In fact, single-cell RNA-seq revealed transcripts encoding the dopamine receptors *Dop1R1, Dop1R2, Dop2R* and/or *DopEcR* in DNT-2+ neurons (Croset et al. 2018). Using reporters, we found that Dop2R is present in DNT-2A neurons (Figure 1F, Figure S1B), but not Dop1R2 (Figure S1E). Altogether, these data showed that DNT-2A neurons are glutamatergic neurons that could receive dopaminergic input both at PRW and SMP. DNT-2 functions via Toll-6 and Kek-6 receptors, and *Toll-6* is expressed in DANs (McIlroy et al. 2013).

**Figure 1.**
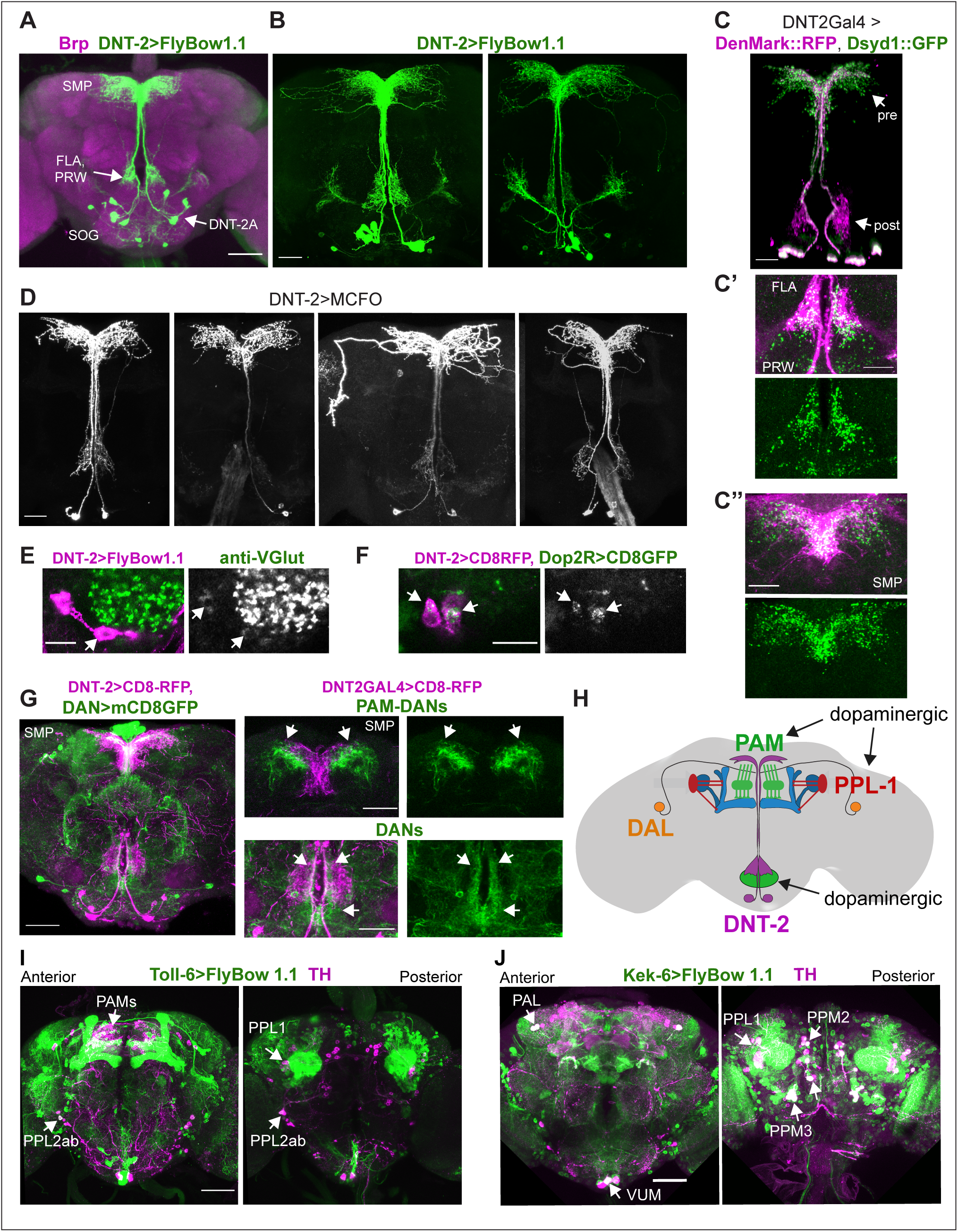
Neurons expressing *DNT-2* and its receptors *Toll-6* and *kek-6* in the adult brain. **(A, B)** *DNT-2A* expressing neurons (*DNT-2>FlyBow1.1* in green; anti-Brp in magenta) have cell bodies in SOG and project to FLA/PRW and SMP. **(C, C’, C’’)** Pre-synaptic (green) and post-synaptic (magenta) terminals of DNT-2A neurons seen with *DNT-2>DenMark::RFP, Dsyd1::GFP*, higher magnification in **(C’,C”),** different specimens from (C). DNT-2A projections at SMP and PRW have both pre- and post-synaptic sites. **(D)** Single neuron DNT-2A>MCFO clones. **(E)** DNT-2A neurons have the vesicular glutamate transporter vGlut (arrows). **(F)** Colocalization between *Dop2RLex>AopCD8-GFP* and *DNT2Gal4>UASCD8-RFP* in cell bodies of DNT-2A neurons (arrows). **(G)** Terminals of dopaminergic neurons (*TH>mCD8GFP*) abut and overlap those of DNT-2A neurons (*DNT2>CD8-RFP*, magenta), arrows; magnified projections on the right. **(H)** Illustration of neurons expressing DNT-2 (magenta) and KCs, DAN PAM and PPL1, and DAL neurons **(I)** *Toll-6>FlyBow1.1* is expressed in Kenyon cells, PPL1, PPL2 and PAM DANs, as revealed by co-localization with anti-TH. **(J)** *kek-6>FlyBow1.1* co-localises with TH in MB vertical lobes, dopaminergic PALs, VUMs, PPL1, PPM2 and PPM3. SMP: superior medial protocerebrum, PRW: Prow, FLA: Flange, SOG: sub-aesophageal ganglion. Scale bars: (A, H, I) 50µm; (B,C,C”, D,G) 30µm (C’, F,) 25µm.

To identify the cells expressing *Toll-6* and *kek-6* and explore further their link to the dopaminergic system, we used *Toll-6Gal4 (Li et al. 2020)* and *kek-6Gal4* (Ulian-Benitez et al. 2017) to drive expression of membrane-tethered *FlyBbow1.1* and assessed their expression throughout the brain. Using anti-Tyrosine Hydroxilase (TH) - the enzyme that catalyses the penultimate step in dopamine synthesis - to visualise DANs, we found that Toll-6+ neurons included dopaminergic neurons from the PAMs, PPL1 and PPL2 clusters (Figure 1I, Figure S2D,F Table S1), whilst Kek-6+ neurons included PAM, PAL, PPL1, PPM2 and PPM3 dopaminergic clusters (Figure 1J, Figure S2B,E,F Table S1). DNT-2 can also bind various Tolls and Keks promiscuously (McIlroy et al. 2013, Foldi et al. 2017) and other *Tolls* are also expressed in the dopaminergic system: PAMs express multiple *Toll* receptors (Figure S2F) and all PPL1s express at least one *Toll* (Figure S2F). Using MCFO clones revealed that both *Toll-6* and *kek-6* are also expressed in Kenyon cells (Li et al. 2020) (Figure S2I,J, Table S1), DAL neurons (Figure S2G,H, Table S1) and MBONs (Figure S2A-C). In summary, *Toll-6* and *kek-6* are expressed in DANs, DAL, Kenyon cells and MBONs (Figure 1H). These cells belong to a circuit required for associative learning, long-term memory and behavioural output, and DANs are also required for locomotion (Riemensperger et al. 2011, Chen et al. 2012, Aso et al. 2014b, Boto et al. 2014, Adel and Griffith 2021, Huang et al. 2024). Altogether, our data showed that DNT-2A neurons are glutamatergic neurons that could receive dopamine as they contacted DANs and expressed the *Dop2R* receptor, and that in turn DANs expressed the DNT-2 receptors *Toll-6* and *kek-6*, and therefore could respond to DNT-2. These data suggested that there could be bidirectional connectivity between DNT-2A neurons and DANs, which we explored below.

### Bidirectional connectivity between DNT-2A neurons and DANs

To verify the connectivity of DNT-2A neurons with DANs, we used various genetic tools. To identify DNT-2A output neurons, we used TransTango (Talay et al. 2017) (Figure 2A and Figure S3). DNT-2A RFP+ outputs included a subset of MB α’β’ lobes, αβ Kenyon cells, tip of MB β’2, DAL neurons, dorsal fan-shaped body layer and possibly PAM or other dopaminergic neurons (Figure 2A, Figure S3). Consistently, these DNT-2A output neurons express *Toll-6* and *kek-6* (Supplementary Table S1). To identify DNT-2A input neurons, we used BAcTrace (Cachero et al. 2020). This identified PAM-DAN inputs at SMP (Figure 2B). Altogether, these data showed that DNT-2A neurons receive dopaminergic neuromodulatory inputs, their outputs include MB Kenyon cells, DAL neurons and possibly DANs, and DNT-2 arborisations at SMP are bidirectional.

**Figure 2.**
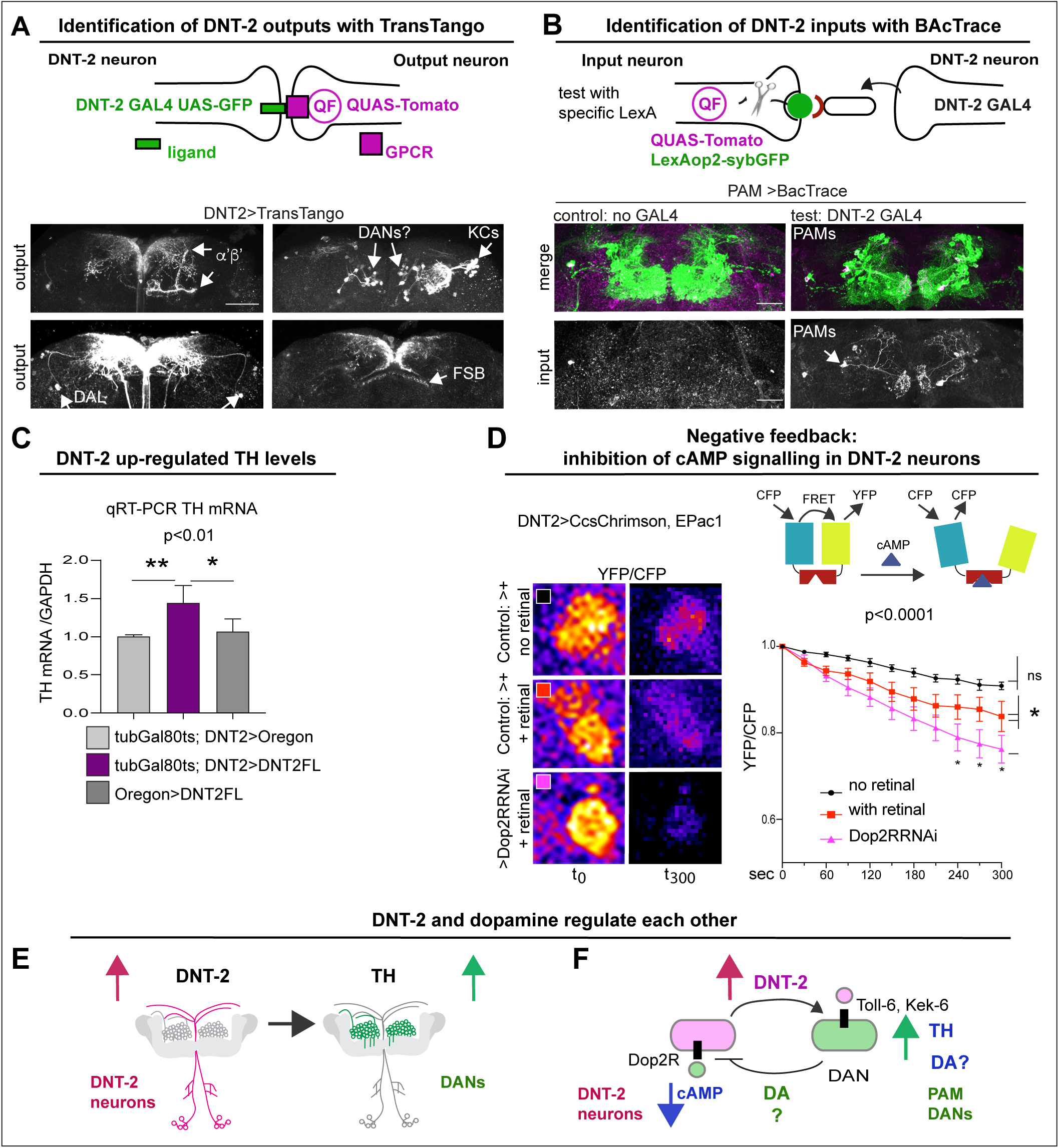
DNT-2 neurons are functionally connected to dopaminergic neurons. **(A)** TransTango revealed out-puts of DNT-2 neurons. All neurons express *TransTango* and expression of the TransTango ligand in DNT-2 neurons identified DNT-2 out-puts with Tomato (anti-DsRed). TransTango identified as DNT-2 outputs KC α’β’ MB lobes (anterior brain, arrow, **top left**); Kenyon and possibly DAN cell bodies (posterior, arrows **top right**); DAL neurons **(bottom left,** arrows) and the dorsal layer of the fan shaped body **(bottom right,** arrow). See also Supplementary Figure S3 for further controls. **(B)** BAcTrace tests connectivity to a candidate neuron input visualised with *LexAop>sybGFP* by driving the expression of a ligand from *DNT-2GAL4* that will activate *QUASTomato* in the candidate input neuron (Cachero et al. 2020). Candidate PAM neurons visualised at SMP with GFP (green): Control: *R58E02LexA>BAcTrace 806*, no *GAL4*. Test: *R58E02LexA, DNT-2GAL4>BAcTrace 806* revealed PAMs are inputs of DNT-2A neurons at SMP (Tomato, bottom). Magenta shows QUAS-Tomato. **(C)** qRT-PCR showing that TH mRNA levels increased with DNT-2 over-expression at 30°C *(tubGAL80^ts^, DNT-2>DNT-2FL).* One way ANOVA, P=0.0085; post-doc Dunnett’s multiple comparison test**. (D)** FRET cAMP probe Epac1 revealed that *DNT-2>Dop2R-RNAi* knockdown decreased YFP/CFP ratio over time in DNT-2A neurons, meaning that cAMP levels increased. Two-way ANOVA, genotype factor p<0.0001, time factor p<0.0001; post-doc Dunnett’s. **(E,F)** Summary: DNT-2 neurons and DANs are functionally connected and modulate each other. **(E)** DNT-2 can induce *TH* expression in DANs; **(F)** this is followed by negative feedback from DANs to DNT-2 neurons (question marks indicate inferences). TH: tyrosine hydroxylase. Scale bars: (A) 50µm; (B) 30µm; (D) 20µm. P values over graphs in (C) refer to group analyses; stars indicate multiple comparisons tests. *p<0.05,**p<0.01, ***p<0.001. For sample sizes and further statistical details, see Supplementary Table S2.

To further test the relationship between DNT-2A neurons and DANs, we reasoned that stimulating DANs would provoke either release or production of dopamine. So, we asked whether increasing DNT-2 levels in DNT-2 neurons could influence dopamine levels. For this, we over-express *DNT-2* in full-length form (i.e. *DNT-2FL*), as it enables to investigate non-autonomous functions of DNT-2 (Ulian-Benitez et al. 2017). Importantly, DNT-2FL is spontaneously cleaved into the mature form (McIlroy et al. 2013, Foldi et al. 2017) (see discussion). Thus, we over-expressed *DNT-2FL* in DNT-2 neurons and asked whether this affected dopamine production, using mRNA levels for *TH* as readout. Using quantitative real-time PCR (qRT-PCR) we found that over-expressing *DNT2-FL* in DNT-2 neurons in adult flies increased *TH* mRNA levels in fly heads (Figure 2C). This showed that DNT-2 could stimulate dopamine production.

Next, we wondered whether in turn DNT-2A neurons, that express *Dop2R*, could be modulated by dopamine. Binding of dopamine to D2-like Dop2R (also known as DD2R) inhibits adenylyl-cyclase, decreasing cAMP levels (Hearn et al. 2002, Neve et al. 2004). Thus, we asked whether DNT-2A neurons received dopamine and signal via Dop2R. Genetic restrictions did not allow us to activate PAMs and test DNT-2 neurons, so we activated DNT-2 neurons and tested whether *Dop2R* knock-down would increase cAMP levels. We used the FRET-based cAMP sensor, Epac1-camps-50A (Shafer et al. 2008). When Epac1-camps-50A binds cAMP, FRET is lost, resulting in decreased YFP/CFP ratio over time. Indeed, *Dop2R* RNAi knock-down in DNT-2A neurons significantly increased cAMP levels (Figure 2D), demonstrating that normally Dop2R inhibits cAMP signalling in DNT-2A cells. Importantly, this result meant that in controls, activating DNT-2A neurons caused dopamine release from DANs that then bound Dop2R to inhibit adenylyl-cyclase in DNT-2A neurons; this inhibition was prevented with *Dop2R* RNAi knock-down. Altogether, this shows that DNT-2 up-regulated TH levels (Figure 2E), and presumably via dopamine release, this inhibited cAMP in DNT-2A neurons (Figure 2F).

In summary, DNT-2A neurons are connected to DANs, DAL and MB Kenyon cells, all of which express DNT-2 receptors *Toll-6* and *kek-6* and belong to a dopaminergic as well as associative learning and memory circuit. Furthermore, DNT-2A and PAM neurons form bidirectional connectivity. Finally, DNT-2 and dopamine regulate each other: DNT-2 increased dopamine levels (Figure 2E), and in turn dopamine via Dop2R inhibited cAMP signalling in DNT-2A neurons (Figure 2F). That is, an amplification was followed by negative feedback. This suggested that a dysregulation in this feedback loop could have consequences for dopamine-dependent behaviours and for circuit remodelling by the DNT-2 growth factor.

### DNT-2 and Toll-6 maintain survival of PAM dopaminergic neurons in the adult brain

Above we showed that DNT-2 and PAM dopaminergic neurons are connected, so we next asked whether loss of function for *DNT-2* or *Toll-6* would affect PAMs. In wild-type flies, PAM-DAN number can vary between 220-250 cells per *Drosophila* brain, making them ideal to investigate changes in cell number (Liu et al. 2012). Maintenance of neuronal survival is a manifestation of structural brain plasticity in mammals, where it depends on the activity-dependent release of the neurotrophin BDNF (Lu et al. 2005, Wang et al. 2022). Importantly, cell number can also change in the adult fly, as neuronal activity can induce neurogenesis via Toll-2, whereas DANs are lost in neurodegeneration models (Feany and Bender 2000, Li et al. 2020). Thus, we asked whether DNT-2 influences PAM-DAN number in the adult brain. We used *THGal4; R58E02Gal4* to visualise nuclear Histone-YFP in DANs (Figure 3A) and counted automatically YFP+ PAMs using a purposely modified DeadEasy plug-in developed for the adult fly brain (Li et al 2020). DeadEasy plug-ins were developed and used before to count cells labelled with sparsely distributed nuclear markers in embryos (Zhu et al. 2008, Forero et al. 2009, Forero et al. 2010a, Forero et al. 2010b, McIlroy et al. 2013), larvae (Kato et al. 2011, Forero et al. 2012, Losada-Perez et al. 2016) and adult (Li et al. 2020) *Drosophila* brains. Here we show that *DNT2^37^/DNT2^18^* mutant adult brains had fewer PAMs than controls (Figure 3B). Similarly, *Toll-6* RNAi knock-down in DANs also decreased PAM neuron number (Figure 3C). DAN loss was confirmed with anti-TH antibodies and counted manually, as there were fewer TH+ PAMs in *DNT2^37^/DNT2^18^*mutants (Figure 3D). Importantly, PAM cell loss was rescued by over-expressing activated *Toll-6^CY^*in DANs in *DNT-2* mutants (Figure 3D). Altogether, these data showed that DNT-2 functions via Toll-6 to maintain PAM neuron survival.

**Figure 3.**
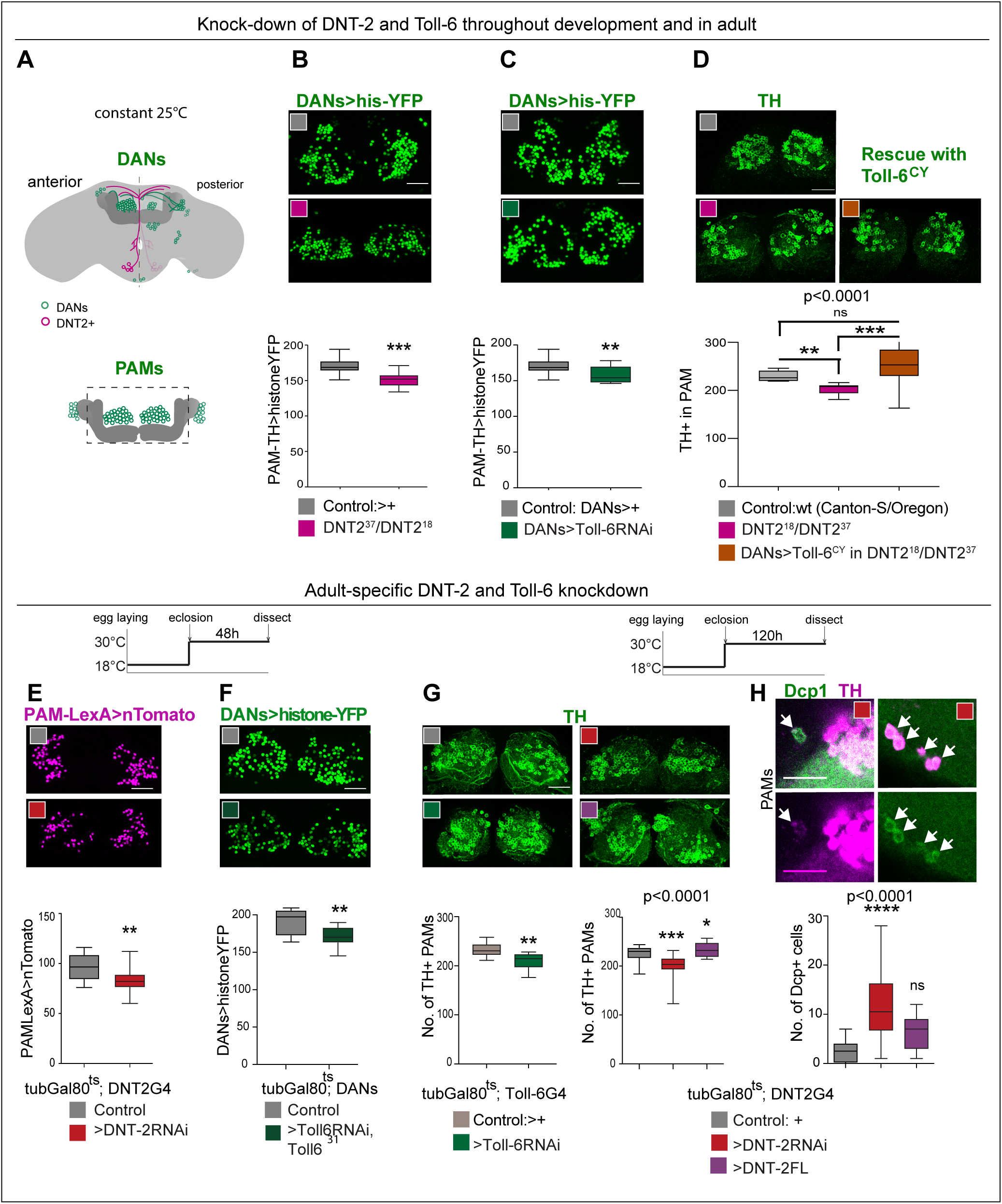
DNT-2 and Toll-6 maintain PAM neuron survival in the developing and adult brain. **(A)** Illustration of PAM neuronal cell bodies and experimental temporal profile. DANs are shown in green, DNT-2A neurons in magenta and MB in dark grey. The left hemisphere shows the anterior brain with PAL and PAM DAN neurons (green) and DNT-2 neurons (magenta); the right shows the posterior brain, with the calyx and other DAN neurons (PPM1, PPM2, PPM3, PPL1, PPL2, green). **(B-D)** Fruit-flies were kept constantly at 25°C, from development to adult. Analyses done in adult brains**. (B)** *DNT-2^37^/DNT-2^18^* mutants had fewer histone-YFP+ labelled PAM neurons (*THGAL4, R58E02-GAL4>hisYFP).* Un-paired Student t-test. **(C)** *Toll-6* RNAi knock-down in all DANs (*THGAL4, R58E02-GAL4>hisYFP, Toll-6 RNAi)* reduced Histone-YFP+ labelled PAM cell number. Un-paired Student t-test. **(D)** *DNT-2^37^/DNT-2^18^* mutants had fewer PAMs stained with anti-TH antibodies. Over-expressing *Toll-6^CY^* in DANs (*THGAL4, R58E02 GAL4>hisYFP, Toll-6^CY^*) rescued TH+ PAM neurons in *DNT-2^37^/DNT-2^18^* mutants, demonstrating that DNT-2 functions via Toll-6 to maintain PAM cell survival. Welch ANOVA p<0.0001, post-hoc Dunnett test. **(E-H)** Adult specific restricted over-expression or knock-down at 30°C, using the temperature sensitive GAL4 repressor *tubGAL80^ts^*. **(E)** Adult specific *DNT-2* RNAi knockdown in DNT-2 neurons decreased Tomato+ PAM cell number (*tubGAL80^ts^, R58E02-LexA, DNT-2 GAL4>LexAOP-Tomato, UAS DNT-2-RNAi*). Un-paired Student t-test, p= 0.005. **(F)** Adult specific *Toll-6* RNAi knock-down in *Toll-6^31^* heterozygous mutant flies in DANs, reduced Histone-YFP+ PAM cell number (*tubGAL80^ts^; THGAL4, R58E02-GAL4>hisYFP, Toll-6 RNAi/Toll6^31^*). Un-paired Student t-test. **(G)** PAMs were visualised with anti-TH. Left: *tubGAL80^ts^, Toll-6>Toll-6-RNAi* knock-down decreased TH+ PAM cell number. Un-paired Student t-test. Right: *tubGAL80^ts^, DNT-2>DNT-2-RNAi* knock-down decreased TH+ PAM cell number, whereas *DNT-2FL* over-expression increased PAM cell number. Kruskal-Wallis ANOVA, p=0.0001, post-hoc Dunn’s test. **(H)** Adult-specific *tubGAL80^ts^, DNT-2>DNT-2RNAi* knock-down increased the number of apoptotic cells in the brain labelled with anti-DCP-1. Dcp-1+ cells co-localise with anti-TH at least in PAM clusters. One Way ANOVA, p<0.0001, post-doc Bonferroni’s multiple comparisons test. DANs>histone-YFP: all dopaminergic neurons expressing histone-YFP, genotype: *THGal4 R58E02Gal4>UAS-histoneYFP*. *PAMsLexA>tomato*: restricted to PAM DANs: *R58E02LexA>LexAop-nlstdTomato.* Controls: *GAL4* drivers crossed to wild-type Canton-S. Scale bars: (B-G) 30µm; (H) 20µm. Graphs show box-plots around the median. P values over graphs in (D, G right, H) refer to group analyses; stars indicate multiple comparisons tests. *p<0.05, **p<0.01, ***p<0.001. For further genotypes, sample sizes and statistical details, see Supplementary Table S2.

To ask whether DNT-2 could regulate dopaminergic neuron number specifically in the adult brain, we used *tubGal80^ts^* to conditionally knock-down gene expression in the adult. PAMs were visualised with either *R58E02LexA>LexAop-nls-tdTomato* or *THGal4; R58E02Gal4* >*histone-YFP* and counted automatically. Adult specific *DNT-2* RNAi knock-down decreased Tomato+ PAM cell number (Figure 3E). Similarly, RNAi *Toll-6* knock-down in DANs also decreased PAM neuron number (Figure 3F). Furthermore, knock-down of either *Toll-6* or *DNT-2* in the adult brain caused loss of PAM neurons visualised with anti-TH antibodies and counted manually (Figure 3G). Cell loss was due to cell death, as adult specific *DNT-2 RNAi* knock-down increased the number of apoptotic cells labelled with anti-*Drosophila* Cleave caspase-1 (DCP-1) antibodies compared to controls, including Dcp-1+ PAMs and other TH+ cells (Figure 3H). Dcp-1+ cells also included TH-negative cells, consistent with the expression of *Toll-6* and *kek-6* also in other cell types. By contrast, *DNT-2FL* over-expression in DNT2 neurons did not alter the incidence of apoptosis (Figure 3G), consistently with the fact that DNT-2FL spontaneously cleaves into the mature form (McIlroy et al. 2013, Foldi et al. 2017). Instead, and importantly, over-expression of DNT-2FL increased PAM cell number (Figure 3G). Thus, *DNT-2* and *Toll-6* knock-down specifically in the adult brain induced apoptosis and PAM-neuron loss, whereas DNT-2 gain of function increased PAM cell number.

Altogether, these data showed that PAM cell number is plastic, sustained PAM neuron survival in development and in the adult brain depends on DNT-2 and Toll-6, and a reduction in their levels causes DAN cell loss, characteristic of neurodegeneration.

### DNT-2 and its receptors are required for arborisations and synapse formation

We next asked whether DNT-2, Toll-6 and Kek-6 could influence dendritic and axonal arbours and synapses of dopaminergic neurons (Figure 4A). Visualising the pre-synaptic reporter Synaptotagmin-GFP (Syt-GFP) in all DANs, we found that *DNT-2^18^/DNT-2^37^*mutants completely lacked DAN synapses in the MB β,β’ and γ lobes (Figure 4B). Interestingly, DAN connections at α,α’ lobes were not affected (Figure 4B). This means that DNT-2 is required for synaptogenesis and connectivity of PAMs to MB β,β’ and γ lobes.

**Figure 4.**
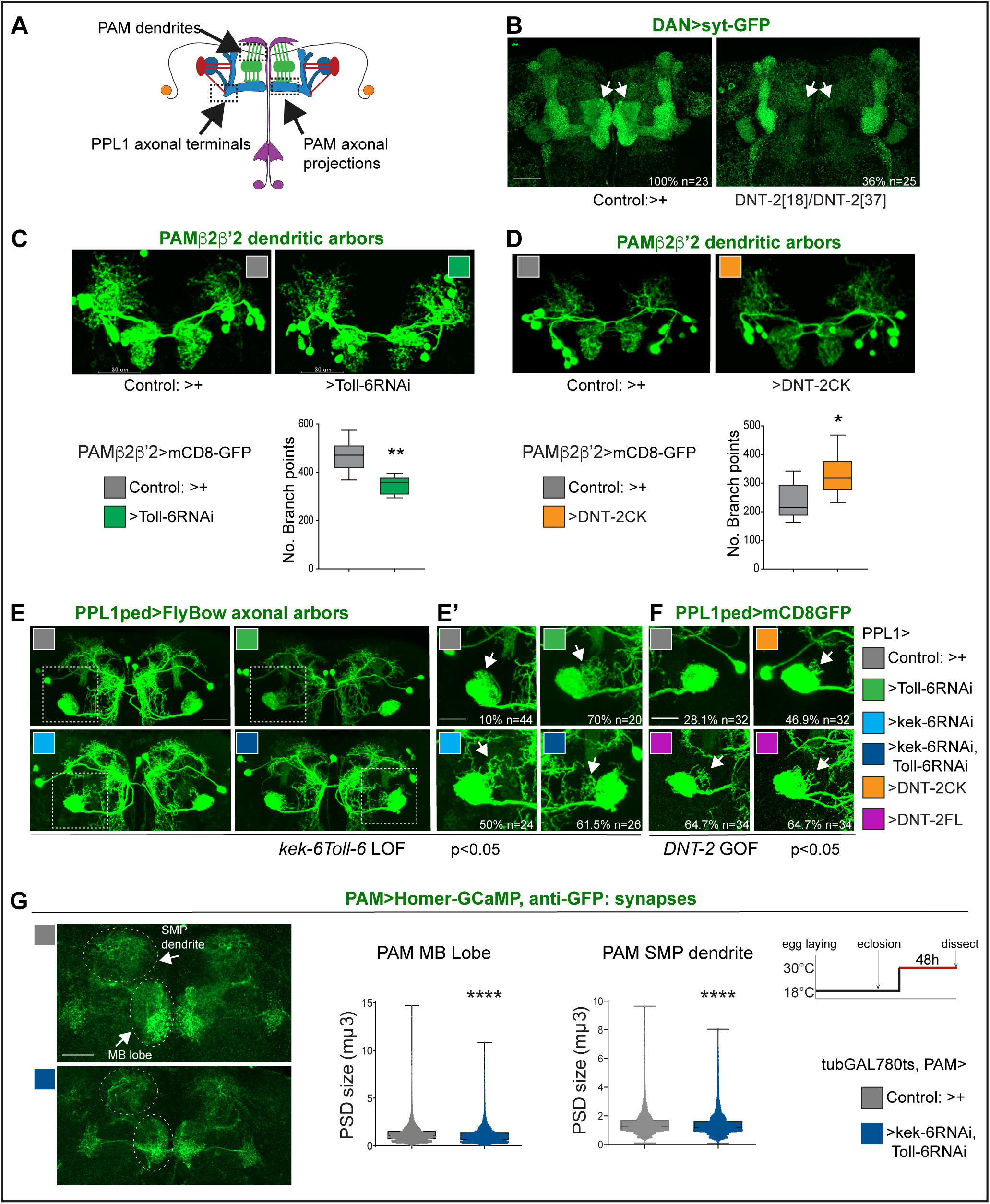
DNT-2, Toll-6 and Kek-6 are required for arborisations and synapse formation. **(A)** Illustration showing the ROIs (dashed lines) used for the analyses, corresponding to dendrites and axonal endings of PAMs and axonal terminals of PPL1ped neurons. **(B-F)** Fruit-flies were kept constantly at 25°C. **(B)** Complete loss of DAN synapses *(TH, R58E02>sytGFP)* onto α,α MB lobe in *DNT-2* null mutants. **(C)** *Toll-6* RNAi knockdown in PAM β2β’2 neurons (*MB301B>FB1.1, Toll-6-RNAi*) decreased dendrite complexity. Un-paired Student t-test. **(D)** Over-expression of cleaved *DNT-2CK* in PAM β2β’2 neurons *(MB301B>CD8-GFP, DNT-2CK)* increased dendrite complexity. Un-paired Student t-test. **(E,E’,F)** PPL1ped axonal misrouting was visualised with *split-GAL4 MB320CGal4>FlyBow1.1.* Images show PPL1-γped neurons and some PPL1-α2α’2. **(D,D’)** RNAi knock-down of *Toll-6, kek-6* or both (e.g. *MB320CGal4>FlyBow1.1, Toll-6RNAi)* in PPL1-γped neurons caused axonal terminal misrouting (arrows). **(E’)** Higher magnification of **(E,** dotted squares). Chi-square for group analysis: p = 0.0224, and multiple comparisons Bonferroni correction control vs *Toll-6RNAi* p<0.01; vs *kek-6RNAi* p<0.05; vs *Toll-6RNAi kek-6RNAi* p<0.01, see Table S2**. (F)** PPL1 misrouting was also induced by over-expressed *DNT-2CK* or *DNT-2FL* (e.g. *MB320CGal4>FlyBow1.1, DNT-2FL)*. Chi-square for group analysis: p<0.05, Bonferroni correction control vs *DNT-2CK* ns, vs *DNT-2FL* *p<0.05, see Table S2. **(G)** Adult-specific *Toll-6 kek-6 RNAi* knockdown in PAM neurons decreased size of post-synaptic density sites (*PAM>Homer-GCaMP* and anti-GFP antibodies). Temperature regime shown on the right. **(C,D)** Graphs show box-plots around the median; **(G)** are box-plots with dot plots. (C,D,G) *p<0.05, **p<0.01, ****p<0.0001. Scale bars: (B,C,D,E) 30µm; (E’,F,G) 20µm. For genotypes, sample sizes and statistical details, see Supplementary Table S2.

PAM-β2β’2 neuron dendrites overlap axonal DNT2 projections. *Toll-6* RNAi knock-down in PAM - β2β’2 (with split-GAL4 *MB301BGal4* (Aso et al. 2014a)), reduced dendrite complexity (Figure 4C). To test whether DNT-2 could alter these dendrites, we over-expressed mature *DNT-2CK*. DNT-2CK is not secreted (from transfected S2 cells), but it is functional in vivo (Zhu et al. 2008, Foldi et al. 2017, Ulian-Benitez et al. 2017). Importantly, over-expressed *DNT-2CK* functions cell-autonomously whereas *DNT-2FL* functions also non-autonomously, but they have similar effects (Zhu et al. 2008, Foldi et al. 2017, Ulian-Benitez et al. 2017). Over-expression of *DNT-2CK* in PAM-β2β’2 increased dendrite arbour complexity (Figure 4D). Thus, DNT-2 and its receptor Toll-6 are required for dendrite growth and complexity in PAM neurons.

To ask whether DNT-2 could affect axonal terminals, we tested PPL1 axons. PPL1-γ1-pedc neurons have a key function in long-term memory gating (Aso et al. 2012, Placais et al. 2012, Aso et al. 2014a, Boto et al. 2020, Huang et al. 2024) and express both *Toll-6* and *kek-6* (Table S1). Using split-GAL4 line M*B320C-Gal4* to visualise PPL1 axonal arbours, RNAi knock-down of either *Toll-6*, *kek-6* or both together caused axonal misrouting away from the mushroom body peduncle (Figure 4E,E’, Chi Square p<0.05, see Table S2). Similarly, *DNT-2FL* over-expression also caused PPL1 misrouting (Figure 4F, Chi-Square p<0.05, see Table S2). Thus, DNT-2, Toll-6 and Kek-6 are required for appropriate targeting and connectivity of PPL1 DAN axons.

To test whether this signalling system was required specifically in the adult brain, we used *tubGAL80^ts^*to knock-down *Toll-6* and *kek-6* with RNAi conditionally in the adult and visualised the effect on synaptogenesis using the post-synaptic reporter Homer-GCaMP and anti-GFP antibodies. Adult-specific *Toll-6 kek-6* RNAi knock-down in PAM neurons did not affect synapse number (not shown), but it decreased post-synaptic density size, both at the MB lobe and at the SMP dendrite (Figure 4G). These data meant that the DNT-2 receptors *Toll-6* and *kek-6* continue to be required in the adult brain for appropriate synaptogenesis.

Altogether, these data showed that DNT-2, Toll-6 and Kek-6 are required for dendrite branching, axonal targeting and synapse formation. The shared phenotypes from altering the levels of *DNT-2* and *Toll-6 kek-6* in arborisations and synapse formation, support their joint function in these contexts. Importantly, these findings showed that connectivity of PAM and PPL1 dopaminergic neurons depend on DNT-2, Toll-6 and Kek-6.

### DNT-2 neuron activation and *DNT-2* over-expression induced synapse formation in target PAM dopaminergic neurons

The above data showed that DNT-2, Toll-6 and Kek-6 are required for DAN cell survival, arborisations and synaptogenesis in development and adults. This meant that the dopaminergic circuit remains plastic in adult flies, consistently with their functional plasticity (Boto et al. 2014). Thus, we wondered whether neuronal activity could also induce remodelling in PAM neurons. In mammals, neuronal activity induces translation, release and cleavage of BDNF, and BDNF drives synaptogenesis (Poo 2001, Lu et al. 2005, Lu et al. 2013, Wang et al. 2022). Thus, we first asked whether neuronal activity could influence DNT-2 levels or function. We visualised tagged DNT-2FL-GFP in adult brains, activated DNT-2 neurons with TrpA1 at 30°C, and found that DNT-2 neuron activation increased the number of DNT-2-GFP vesicles produced (Supplementary Figure S4A). Furthermore, neuronal activity also facilitated cleavage of DNT-2 into its mature form. In western blots from brains over-expressing *DNT-2FL-GFP*, the levels of full-length DNT-2FL-GFP were reduced following neuronal activation and the cleaved DNT-2CK-GFP form was most abundant (Supplementary Figure S4B). These findings meant that, like mammalian BDNF, also DNT-2 can be influenced by activity.

Thus, we asked whether neuronal activity and DNT-2 could influence synapse formation. We first tested DNT-2 neurons. Activating DNT-2 neurons altered DNT-2 axonal arbours (Figure 5A) and it increased Homer-GFP+ synapse number in the DNT-2 SMP arbour (Figure 5B and Figure S5). Next, as DNT-2 and PAMs form bidirectional connexions at SMP (Figure 1, 2), we asked whether activating DNT-2 neurons could affect target PAM neurons. To manipulate DNT-2 neurons and visualise PAM neurons concomitantly, we combined *DNT-2GAL4* with the *PAM-LexAOP* driver. However, there were no available *LexA/OP* post-synaptic reporters, so we used the pre-synaptic *LexAOP-Syt-GCaM*P reporter instead, which labels Synaptotagmin (Syt), and GFP antibodies. Activating DNT-2 neurons with TrpA1 increased the number of Syt+ synapses at the PAM SMP arbour (Figure 5C) and reduced their size (Figure 5C). This was consistent with the increase in Homer-GFP+ PSD number in stimulated DNT-2 neurons (Figure 5B). Neuronal activity can induce ghost boutons, immature synapses that are later eliminated (Fuentes-Medel et al. 2009). Here, the coincidence of increased pre-synaptic Syt-GFP from PAMs and post-synaptic Homer-GFP from DNT-2 neurons at SMP suggests that newly formed synapses could be stable. PAM neurons also send an arborisation at the MB β, β’, γ lobes, but DNT-2 neuron activation did not affect synapse number nor size there (Figure 5C). These data showed that activating DNT-2 neurons induced synapse formation at the SMP connection with PAMs.

**Figure 5.**
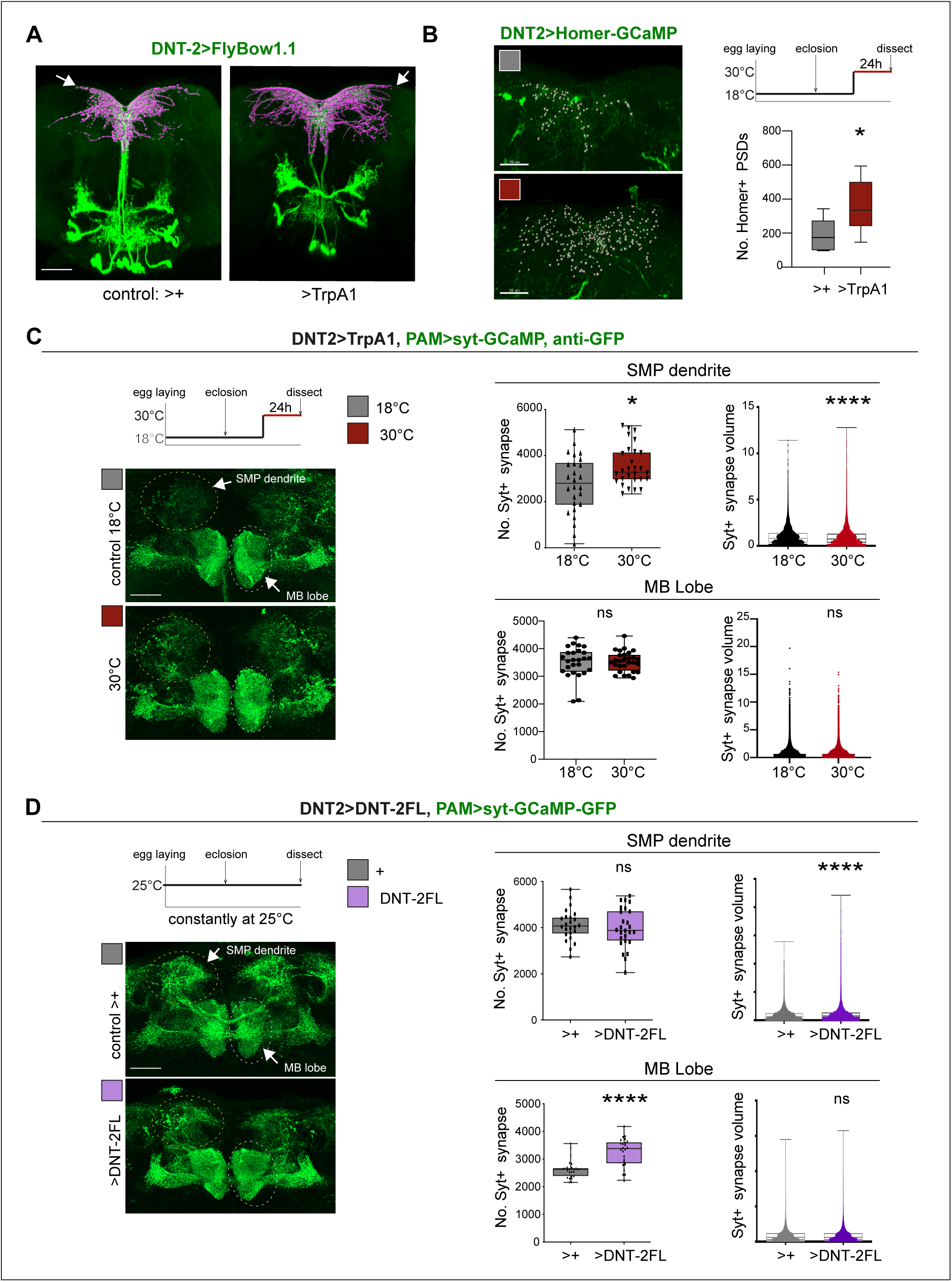
DNT-2 neuron activation and DNT-2 over-expression induced synaptogenesis. **(A)** Thermo-activation of DNT-2 neurons at 30°C altered DNT-2 arborisations, here showing an example with smaller dendrites and enlarged axonal arbours (magenta shows 3D-rendering of axonal arborisation done with Imaris, merged with raw image in green). (Genotype: *DNT-2>FlyBow1.1, TrpA1*). **(B)** Thermogenetic activation of DNT-2 neurons increased the number of Homer+ PSDs in DNT-2 neurons, at SMP (*DNT-2>Homer-GCAMP3, TrpA1*, anti-GFP). Test at 30°C 24h: Unpaired Student t test. See Table S2. **(C)** Thermogenetic activation of DNT-2 neurons induced synaptogenesis in PAM target neurons at SMP, but not at MB lobe. (Genotype: *PAM(R58E02)LexA/LexAOP-sytGCaMP; DNT-2GAL4/UASTrpA1*). At SMP: No. Syt+ synapses: Mann Whitney-U; Syt+ synapse volume: Mann Whitney-U. At MB lobe: No. Syt+ synapses: Unpaired Student t ns; Syt+ synapse volume: Mann Whitney-U ns. **(D)** Over-expression of *DNT-2FL* in DNT-2 neurons increased synapse volume at SMP dendrite and induced synaptogenesis at MB lobe. (Genotype: *PAM(R58E02)LexA/LexAopSytGcaMP6; DNT-2Gal4/UAS-DNT-2FL*). At SMP: No. Syt+ synapses: Unpaired Student t ns. Syt+ synapse volume: Mann Whitney-U. At MB lobe: No. Syt+ synapses: Unpaired Student t; Syt+ synapse volume: Mann Whitney-U ns. Graphs show box-plots around the median, except for PSD volume data that are dot plots. *p<0.05, ****p<0.0001; ns: not significantly different from control. Scale bars: 30µm. For Genotypes, sample sizes, p values and other statistical details, see Supplementary Table S2.

Finally, we asked whether, like activity, DNT-2FL could also drive synaptogenesis. We over-expressed *DNT-2FL* in DNT-2 neurons and visualised the effect in PAM neurons. Over-expression of *DNT-2FL* in DNT-2 neurons did not alter Syt+ synapse number at the PAM SMP dendrite, but it increased bouton size (Figure 5D). By contrast, at the MB β, β’ lobe arborisation, over-expressed *DNT-2* did not affect Syt+ bouton size, but it increased the number of output synapses (Figure 5D). This data showed that DNT-2 released from DNT-2 neurons could induce synapse formation in PAM target neurons.

Altogether, these data showed that neuronal activity induced synapse formation, it stimulated production and cleavage of DNT-2, and DNT-2 could induce synapse formation in target neurons.

### Structural plasticity by DNT2 modified dopamine-dependent behaviour

Circuit structural plasticity raises the important question of what effect it could have on brain function, i.e. behaviour. Data above showed that DANs and DNT-2 neurons are functionally connected, that loss of function for *DNT-2* or its receptors caused dopaminergic neuron loss, altered DAN arborisations and caused synapse loss or reduction in size, and that DNT-2 could induce dendrite branching and synaptogenesis, altogether modifying circuit connectivity. To measure the effect of such circuit modifications on brain function, we used dopamine-dependent behaviours as readout.

Startle-induced negative geotaxis (also known as the climbing assay) is commonly used as a measure of locomotor ability and requires dopamine and specifically PAM neuron function (Riemensperger et al. 2013, Sun et al. 2018). We tested the effect of *DNT-2* or *Toll-6* and *kek-6* loss of function in climbing, and both *DNT-2^37^/DNT-2^18^*mutants and flies in which *DNT-2* was knocked-down in DNT-2 neurons in the adult stage had lower climbing ability than controls (Figure 6A). Similarly, when *Toll-6* and *kek-6* were knocked-down with RNAi in the adult using a *Toll-6-* or a *PAM-GAL4* neuron driver, climbing was also reduced (Figure 6B). Importantly, over-expressing activated *Toll-6^CY^*in DANs rescued the locomotion deficits of *DNT-2* mutants, showing that DNT-2 functions via Toll-6 in this context (Figure 6C).

**Figure 6.**
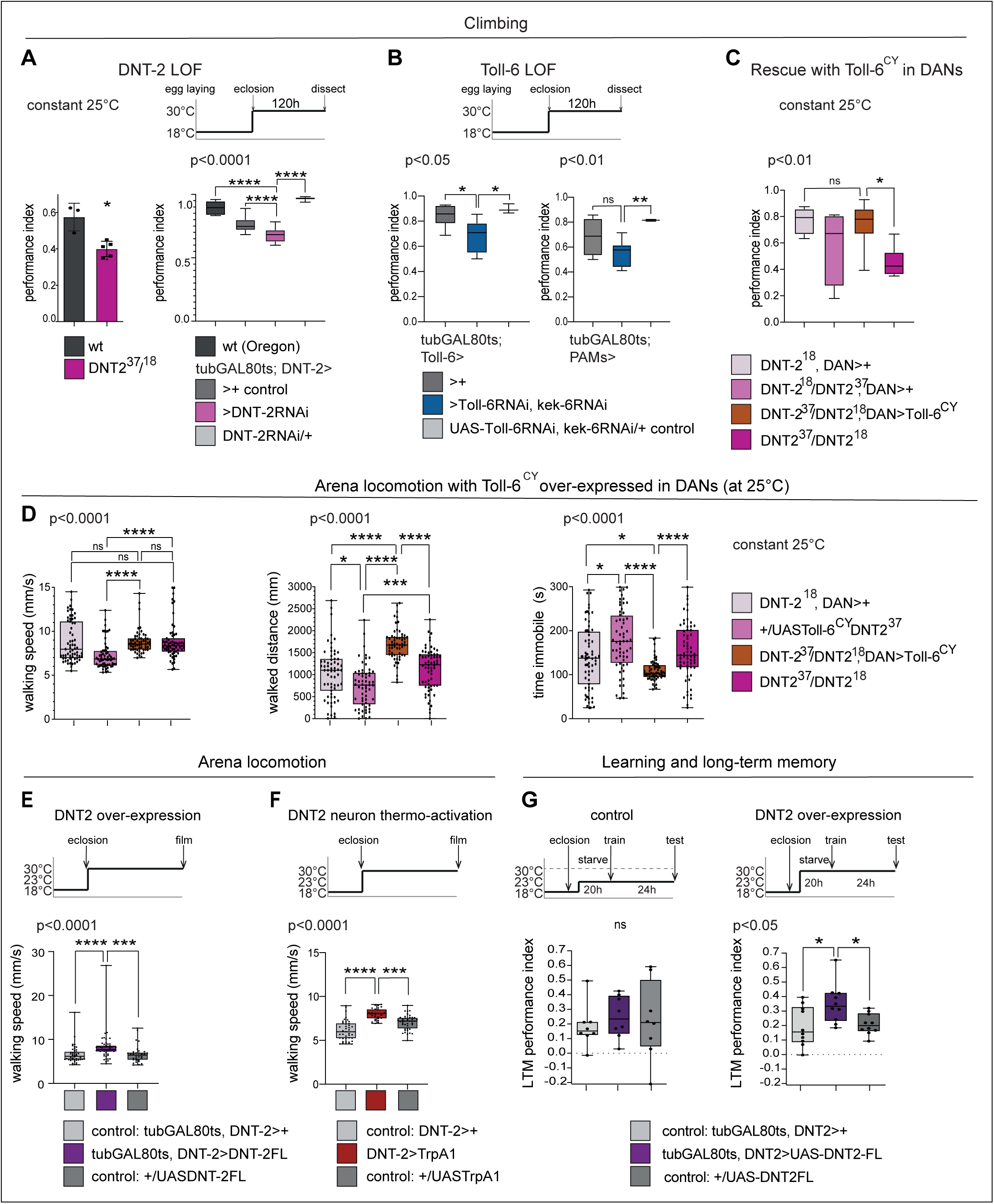
DNT-2-induced circuit plasticity modified dopamine-dependent behaviour. **(A)** *DNT-2* mutants (left, *DNT-2^37^/DNT-2^18^*) and flies with adult-specific RNAi knock-down of *DNT-2* in DNT-2 neurons (right, *tubGAL80^ts^; DNT-2>DNT-2RNAi*), had impaired climbing. Left Mann Whitney U; Right: One Way ANOV, post-hoc Dunnett. **(B)** Adult-specific *Toll-6* and *kek-6* RNAi knock-down in Toll-6 *(tubGAL80^ts^; Toll-6>Toll-6RNAi, kek-6RNAi,* left) or PAM (*tubGAL80^ts^; R58E02>Toll-6RNAi, kek-6RNAi,* right) neurons impaired climbing. Left: Welch ANOVA, post-hoc Dunnett. Right: Welch ANOVA, post-hoc Dunnett. **(C)** The climbing impairment of *DNT-2* mutants could be rescued with the over-expression of *Toll-6^CY^* in dopaminergic neurons (Rescue genotype: UASToll-6^CY^/+; DNT-2^18^*THGAL4 R58E02GAL4/DNT-2^37^).* Welch ANOVA, post-hoc Dunnett. **(D)** Adult specific over-expression of *Toll-6^CY^* in DANs increased locomotion in *DNT-2^37^/DNT-2^18^*mutants. (Test genotype: *UASToll-6^CY^/+; DNT-2^18^THGAL4 R58E02GAL4/DNT-2^37^*). Walking speed: Kruskal Wallis ANOVA, post-hoc Dunn’s. Distance walked: Kruskal Wallis ANOVA, post-hoc Dunn’s. Time spent immobile: Kruskal Wallis ANOVA, post-hoc Dunn’s. **(E)** Adult specific *DNT-2FL* overexpression in DNT-2 neurons increased fruit-fly locomotion speed in an open arena at 30°C (see also Supplementary Figure S6A for further controls). (Test genotype: *tubGAL80^ts^, DNT-2>DNT-2FL*) Kruskal-Wallis, post-hoc Dunn’s test. **(F)** Thermogenetic activation of DNT-2 neurons at 30°C *(DNT-2>TrpA1)* increased fruit-fly locomotion speed. (See also Supplementary Figure S6B for further controls). One Way ANOVA p<0.0001, post-hoc Dunnett’s test**. (G)** Over-expression of *DNT-2-FL* in DNT-2 neurons increased long-term memory. (Test genotype: *tubGAL80^ts^, DNT-2>DNT-2FL*). Left: 23°C controls: One Way ANOVA p=0.8006. Right 30°C: One Way ANOVA, post-hoc Dunnett’s test. Graphs show box-plots around the median, under also dot plots. P values over graphs refer to group analyses; asterisks indicate multiple comparisons tests. *p<0.05, **p<0.01, ***p<0.001, ****p<0.0001; ns: not significantly different from control. For further genotypes, sample sizes, p values and other statistical details, see Supplementary Table S2.

We also tracked freely moving flies in an open arena (Eyjolfsdottir et al. 2014). Interestingly, in that setting, locomotion of homozygous *DNT-2^37^/DNT-2^18^*mutants was similar to that of controls, but over-expression of *Toll-6^CY^*in their DANs increased locomotion as flies walked longer distances and spent less time immobile (Figure 6D). Adult flies over-expressing *DNT2-FL* walked faster (Figure 6E and Figure S5) and so did those where DNT-2 neurons were activated with TrpA1 (Figure 6F and Figure S6), consistently with the fact that neuronal activity increased DNT-2 production (Figure S4A) and that DNT-2FL increased TH levels (Figure 2C). Therefore, increased Toll-6^CY^ levels in DANs increase locomotion and increased DNT-2 levels are sufficient to boost walking speed. Interestingly, both loss and gain of function for *DNT-2* also caused seizures (Figure S7). Thus, dopamine-dependent locomotion is regulated by the function of DNT-2, Toll-6 and Kek-6.

Next, as dopamine is an essential neurotransmitter for learning and memory (Adel and Griffith 2021), we asked whether DNT-2 might influence appetitive olfactory conditioning. Starved flies were trained to associate a sugar reward with an odour (CS+) while another odour was presented without sugar (CS-), and their preference for CS+ versus CS-was measured, 24h after training (Tempel et al. 1983, Krashes and Waddell 2008, 2011). Remarkably, over-expression of *DNT-2FL* in DNT-2 neurons in adults enhanced appetitive long-term memory (Figure 6G), consistent with the positive role of DNT-2 in synaptogenesis that we demonstrated above.

In summary, we have shown that alterations in DNT-2, Toll-6 and Kek-6 levels that caused structural phenotypes in DANs also modified dopamine-dependent behaviours, locomotion and long-term memory.

## DISCUSSION

Our findings indicate that structural plasticity and degeneration in the brain are two manifestations of a shared molecular mechanism that could be modulated by experience. Loss of function for *DNT-2, Toll-6* and *kek-6* caused cell loss, affected arborisations and synaptogenesis in DANs and impaired locomotion; neuronal activity increased DNT-2 production and cleavage and remodelled connecting DNT-2 and PAM synapses; and over-expression of *DNT-2* increased TH levels, PAM cell number, dendrite complexity, and synaptogenesis, and it enhanced locomotion and long-term memory.

It was remarkable to find that the number of dopaminergic neurons in the *Drosophila* adult brain is plastic, and this is functionally relevant as it can influence behaviour. We showed that PAM cell number is variables across individuals, that adult-specific gain of *DNT-2* function increases, whereas loss of *DNT-2* or *Toll-6* function decreases, PAM cell number. Loss of *DNT-2* function in mutants, constant loss of *Toll-6* function in DANS and adult-restricted knock-down of either *DNT-2* (in DNT-2 neurons) or *Toll-6* (in Toll-6 neurons and in DANs) all resulted in DAN cell loss, verified with three distinct reporters, and consistently with the increase in DAN apoptosis. Furthermore, DAN cell loss in *DNT-2* mutants could be rescued by the over-expression of *Toll-6* in DANs. Cell loss was also verified using two reporter types (ie GAL4 based nuclear reporters and cytoplasmic anti-TH antibodies), multiple GAL4 drivers and mutants, and multiple cell counting methods, including automatic cell counting with DeadEasy plug-ins for His-YFP and nls-Tomato (where the signal was of high contrast and sphericity) and software assisted manual cell counting for anti-TH (where the signal is more diffuse and less regular in shape). DeadEasy plug-ins have been used before for reliably counting His-YFP labelled cells in both larval CNS and adult brains, including Kenyon cells (Kato et al. 2011, Forero et al. 2012, Losada-Perez et al. 2016, Li et al. 2020, Harrison et al. 2021). Thus, the finding that loss of *DNT-2* and *Toll-6* function in the adult brain cause dopaminergic neuron loss is robust. Our findings are reminiscent of the increased apoptosis and cell loss in adult brains with *Toll-2* loss of function (Li et al. 2020), and the support of DAN survival by Toll-1 and Toll-7 driven autophagy (Zhang et al. 2024). They are also consistent with a report that loss of function for *DNT-2* or *Toll-6* induced apoptosis in the third instar larval optic lobes (McLaughlin et al. 2019). This did not result in neuronal loss - which was interpreted as due to Toll-6 functions exclusive to glia (McLaughlin et al. 2019) - but instead of testing the optic lobes, neurons of the larval abdominal ventral nerve cord (VNC) were monitored (McLaughlin et al. 2019). In the VNC, Toll-6 and −7 function redundantly and knock-down of both is required to cause neuronal loss in embryos (McIlroy et al. 2013), whereas in L3 larvae and pupae the phenotype is compounded by their pro-apoptotic functions (Foldi et al. 2017). It is crucial to consider that the DNT-Toll signalling system can have distinct cellular outcomes depending on context, cell type and time, i.e. stage (Foldi et al. 2017, Li et al. 2020, Li and Hidalgo 2021). Our work shows that in the adult *Drosophila* brain DAN neurons receive secreted growth factors that maintain cellular integrity, and this impacts behaviour. Consistently with our findings, *Drosophila* models of Parkinson’s Disease reproduce the loss of DANs and locomotion impairment of human patients (Feany and Bender 2000, Riemensperger et al. 2011, Riemensperger et al. 2013, Sun et al. 2018). Dopamine is required for locomotion, associative reward learning and long-term memory (Riemensperger et al. 2011, Riemensperger et al. 2013, Waddell 2013, Sun et al. 2018, Boto et al. 2020, Adel and Griffith 2021). In *Drosophila*, this requires PAM, PPL1 and DAL neurons and their connections to Kenyon Cells and MBONs (Heisenberg 2003, Chen et al. 2012, Placais et al. 2012, Aso et al. 2014b, Placais et al. 2017, Boto et al. 2020, Adel and Griffith 2021, Huang et al. 2024). DNT-2 neurons are connected to all these neuron types, which express *Toll-6* and *kek-6,* and modifying their levels affects locomotion and long-term memory. Altogether, our data demonstrate that structural changes caused by altering DNT-2, Toll-6 and Kek-6 modified dopamine-dependent behaviours, providing a direct link between molecules, structural circuit plasticity and behaviour.

We used neuronal activation as a proxy for experience, but the implication is that experience would similarly drive the structural modification of circuits labelled by neuromodulators. Similar manipulations of activity have previously revealed structural circuit modifications. For example, hyperpolarising olfactory projection neurons increased microglomeruli number, active zone density and post-synaptic site size in the calyx, whereas inhibition of synaptic vesicle release decreased the number of microglomeruli and active zones (Kremer et al. 2010). There is also evidence that experience can modify circuits and behaviour in *Drosophila.* For example, natural exposure to light and dark cycles maintains the structural homeostasis of presynaptic sites in photoreceptor neurons, which breaks down in sustained exposure to light (Sugie et al. 2015). Prolonged odour exposure causes structural reduction at the antennal lobe and at the output pre-synaptic sites in the calyx, and habituation (Devaud et al. 2001, Pech et al. 2015). Similarly, prolonged exposure to CO2 caused a reduction in output responses at the lateral horn and habituation (Sachse et al. 2007). Our findings are also consistent with previous reports of structural plasticity during learning in *Drosophila.* Hypocaloric food promotes structural plasticity in DANs, causing a reduction specifically in connections between DANs and Kenyon cells involved in aversive learning, thus decreasing the memory of the aversive experience (Coban et al. 2024). By contrast, after olfactory conditioning, appetitive long-term memory increased axonal collaterals in projection neurons, and synapse number at Kenyon cell inputs in the calyx (Baltruschat et al. 2021). Our data provide a direct link between a molecular mechanism, synapse formation in a dopaminergic circuit and behavioural performance. Since behaviour is a source of experience, the discovery that a neurotrophin can function with a Toll and a neuromodulator to sculpt circuits provides a mechanistic basis for how experience can shape the brain throughout life.

Importantly, in humans structural brain plasticity (e.g. adult neurogenesis, neuronal survival, neurite growth and synaptogenesis) correlates with anti-depressant treatment, learning, physical exercise and well-being (Cotman and Berchtold 2002, Woollett and Maguire 2011, Castren and Monteggia 2021, Cheng et al. 2022). Conversely, neurite, synapse and cell loss, correlate with ageing, neuroinflammation, psychiatric and neurodegenerative conditions (Holtmaat and Svoboda 2009, Lu et al. 2013, Wohleb et al. 2016, Forrest et al. 2018, Vahid-Ansari and Albert 2021, Wang et al. 2022). Understanding how experience drives the switch between generative and destructive cellular processes shaping the brain is critical to understand brain function, in health and disease. In this context, the mechanism we have discovered could also operate in the human brain. In fact, there is deep evolutionary conservation in DNT-2 vs mammalian NT function (e.g. BDNF), but some details differ. Like mammalian NTs, full-length DNTs/Spz proteins contain a signal peptide, an unstructured pro-domain and an evolutionarily conserved cystine-knot of the NT family (Zhu et al. 2008, Arnot et al. 2010, Foldi et al. 2017). Cleavage of the pro-domain releases the mature cystine-knot (CK). In mammals, full-length NTs have opposite functions to their cleaved forms (e.g. apoptosis vs cell survival, respectively). However, DNT-2FL is cleaved by intracellular furins, and although DNT-2 can be found both in full-length or cleaved forms in vivo, it is most abundantly found cleaved (Zhu et al. 2008, McIlroy et al. 2013, Foldi et al. 2017). As a result, over-expressed *DNT-2FL* does not induce apoptosis and instead it promotes cell survival (Foldi et al. 2017). The same functions are played by over-expressed mature *DNT-2CK* as by *DNT-2FL* (Zhu et al. 2008, Foldi et al. 2017, Ulian-Benitez et al. 2017). In S2 cells, transfected DNT-2CK is not secreted, but when over-expressed in vivo it is functional (Zhu et al. 2008, Foldi et al. 2017, Ulian-Benitez et al. 2017) (and this work). In fact, over-expressed *DNT-2CK* also maintains neuronal survival, connectivity and synaptogenesis (Zhu et al. 2008, Foldi et al. 2017, Ulian-Benitez et al. 2017) (and this work). Similarly, over-expressed mature *spz-1-C106* can rescue the *spz-1*-null mutant phenotype (Hu et al. 2004) and over-expressed *DNT-1CK* can promote neuronal survival, connectivity and rescue the *DNT-1* mutant phenotype (Zhu et al. 2008). Consistently, both DNT-2FL and DNT-2CK have neuro-protective functions promoting cell survival, neurite growth and synaptogenesis (Zhu et al. 2008, Sutcliffe et al. 2013, Foldi et al. 2017, Ulian-Benitez et al. 2017) (and this work). Importantly, over-expressing *DNT-2FL* enables to investigate non-autonomous functions of DNT-2 (Ulian-Benitez et al. 2017). We have shown that *DNT-2FL* can induce synaptogenesis non-autonomously in target neurons. Similarly, DNT-2 is a retrograde factor at the larval NMJ, where transcripts are located post-synaptically in the muscle, DNT-2FL-GFP is taken up from muscle by motoneurons, where it induces synaptogenesis (Sutcliffe et al. 2013, Ulian-Benitez et al. 2017). Importantly, we have shown that neuronal activity increased production of tagged DNT-2-GFP, and its cleavage. In mammals, neuronal activity induces synthesis, release and cleavage of BDNF, leading to neuronal survival, dendrite growth and branching, synaptogenesis and synaptic plasticity (ie LTP)(Poo 2001, Horch and Katz 2002, Lu et al. 2005, Arikkath 2012, Wang et al. 2022). Like BDNF, DNT-2 also induced synaptogenesis and increased bouton size.

It may not always be possible to disentangle primary from compensatory phenotypes, as Hebbian, homeostatic and heterosynaptic plasticity can concur (Forrest et al. 2018, Jenks et al. 2021). Mammalian BDNF increases synapse number, spine size and long-term potentiation (LTP), but it can also regulate homeostatic plasticity and long-term depression (LTD), depending on timing, levels and site of action (Poo 2001, Lu et al. 2005, Wang et al. 2022). In this context, neuronal stimulation of DNT-2 neurons induced synapse formation in PAM neuron SMP dendrites, whereas DNT-2 over-expression from DNT-2 neurons increased synapse size at SMP and synapse number in PAM outputs at the mushroom body lobes. These distinct effects could be due to the combination of plasticity mechanisms and range of action. Neuronal activity can induce localised protein synthesis that facilitates local synaptogenesis and stabilises emerging synapses (Forrest et al. 2018). By contrast, DNT-2 induced signalling via the nucleus can facilitate synaptogenesis at longer distances, in output sites. In any case, synaptic remodelling is the result of concurring forms of activity-dependent plasticity altogether leading to modification in connectivity patterns (Forrest et al. 2018, Jenks et al. 2021). Long-term memory requires synaptogenesis, and in mammals this depends on BDNF, and its role in the protein-synthesis dependent phase of LTP (Poo 2001, Minichiello 2009, Wang et al. 2022). BDNF localised translation, expression and release are induced to enable long-term memory (Poo 2001, Lee et al. 2004, Minichiello 2009, Wang et al. 2022). We have shown that similarly, over-expressed *DNT-2FL* increased both synaptogenesis in sites involved in reward learning, and long-term memory after appetitive conditioning.

The relationship of NTs with dopamine is also conserved. DNT-2 and DAN neurons form bidirectional connectivity that modulates both DNT-2 and dopamine levels. Similarly, mammalian NTs also promote dopamine release and the expression of DA receptors (Blochl and Sirrenberg 1996, Guillin et al. 2001). Furthermore, DAN cell survival is maintained by DNT-2 in *Drosophila,* and similarly DAN cell survival is also maintained by NT signalling in mammals and fish (Hyman et al. 1991, Sahu et al. 2019). Importantly, we showed that activating DNT-2 neurons increased the levels and cleavage of DNT-2, up-regulated DNT-2 increased *TH* expression, and this initial amplification resulted in the inhibition of cAMP signalling via the dopamine receptor Dop2R in DNT-2 neurons. This negative feedback could drive a homeostatic reset of both DNT-2 and dopamine levels, important for normal brain function. In fact, we showed that alterations in DNT-2 levels could cause seizures. Importantly, alterations in both NTs and dopamine underlie many psychiatric disorders and neurodegenerative diseases in humans (Hyman et al. 1991, Guillin et al. 2001, Berton et al. 2006, Forrest et al. 2018, Wang et al. 2022).

We have uncovered a novel mechanism of structural brain plasticity, involving a NT ligand functioning via a Toll and a kinase-less Trk-family receptor in the adult *Drosophila* brain. Toll receptors in the CNS can function via ligand dependent and ligand-independent mechanisms (Anthoney et al. 2018). However, in the context analysed, Toll-6 and Kek-6 function in structural circuit plasticity depends on their ligand DNT-2. This is also consistent with their functions promoting axonal arbour growth, branching and synaptogenesis at the NMJ (Ulian-Benitez et al. 2017). Furthermore, Toll-2 is also neuro-protective in the adult fly brain, and loss of *Toll-2* function caused neurodegeneration and impaired behaviour (Li et al. 2020). There are six *spz/DNT*, nine *Toll* and six *kek* paralogous genes in *Drosophila* (Tauszig et al. 2000, MacLaren et al. 2004, Mandai et al. 2009, Ulian-Benitez et al. 2017), and at least seven *Tolls* and three adaptors are expressed in distinct but overlapping patterns in the brain (Li et al. 2020). Such combinatorial complexity opens the possibility for a fine-tuned regulation of structural circuit plasticity and homeostasis in the brain.

*Drosophila* and mammalian NTs may have evolved to use different receptor types to elicit equivalent cellular outcomes. In fact, in mammals, NTs function via Trk, p75^NTR^ and Sortilin receptors, to activate ERK, PI3K, NFκB, JNK and CaMKII (Lu et al. 2005, Wang et al. 2022). Similarly, DNTs with Tolls and Keks also activate ERK, NFκB, JNK and CaMKII in the *Drosophila* CNS (McIlroy et al. 2013, Foldi et al. 2017, Ulian-Benitez et al. 2017, Anthoney et al. 2018). However, alternatively, NTs may also use further receptors in mammals. Trks have many kinase-less isoforms, and understanding of their function is limited (Fryer et al. 1996, Stoilov et al. 2002). They can function as ligand sinks and dominant negative forms, but they can also function independently of full-length Trks to influence calcium levels, growth cone extension and dendritic growth and are linked to psychiatric disorders, e.g. depression (Ferrer et al. 1999, Yacoubian and Lo 2000, Ohira et al. 2006, Carim-Todd et al. 2009, Ernst et al. 2009, Ohira and Hayashi 2009, Fenner 2012, Tessarollo and Yanpallewar 2022). Like Keks, perhaps kinase-less Trks could regulate brain plasticity vs. degeneration.

A functional relationship between NTs and TLRs could exist also in humans, as in cell culture, human BDNF and NGF can induce signalling from a TLR (Foldi et al. 2017) and NGF also functions in immunity and neuroinflammation (Levi-Montalcini et al. 1996, Hepburn et al. 2014). Importantly, TLRs can regulate cell survival, death and proliferation, neurogenesis, neurite growth and collapse, learning and memory (Okun et al. 2011). They are linked to neuroinflammation, psychiatric disorders, neurodegenerative diseases and stroke (Okun et al. 2011, Figueroa-Hall et al. 2020, Adhikarla et al. 2021). Intriguingly, genome-wide association studies (GWAS) have revealed the involvement of TLRs in various brain conditions and potential links between NTs and TLRs in, for example, major depression (Sharma 2012, Mehta et al. 2018, Chan et al. 2020, Garrett et al. 2021). Importantly, alterations in NT function underlie psychiatric, neurologic and neurodegenerative brain diseases (Lu et al. 2005, Martinowich et al. 2007, Krishnan and Nestler 2008, Lu et al. 2013, Park and Poo 2013, Wohleb et al. 2016, Yang et al. 2020, Casarotto et al. 2021, Wang et al. 2022) and BDNF underlies the plasticity inducing function of anti-depressants (Lu et al. 2013, Casarotto et al. 2021, Wang et al. 2022). It is compelling to find out whether and how these important protein families – NTs, TLRs and kinase-less Trks - interact in the human brain.

## Conclusion

To conclude, we provide a direct link between structural circuit plasticity and behavioural performance, by a novel molecular mechanism. The neurotrophin DNT-2 and its receptors Toll-6 and the kinase-less Trk family Kek-6 are linked to a dopaminergic circuit. Neuronal activity boosts DNT-2, and DNT-2 and dopamine regulate each other homeostatically. Dopamine labels the circuits engaged and DNT-2, a growth factor, with its receptors Toll-6 and Kek-6, drives structural plasticity in these circuits, enhancing dopamine-dependent behavioural performance. These findings mean that DNT-2 is a plasticity factor in the *Drosophila* brain, that could enable experience-dependent behavioural enhancement. Whether NTs can similarly functions with TLRs and Kinase-less Trks remains to be explored. As behaviour is a source of experience, this has profound implications for understanding brain function and health.

## ACKNOWLEDGEMENTS

We thank our lab, Carolina Rezaval and Thomas Riemensperger for comments on the manuscript; Carolina Rezaval, Reinhard Wolf, Martin Heisenberg and Scott Waddell for advice on - Karina Piotrowska for help with - behaviour experiments; Xiufeng Li for help with programming; Serge Birman, Ann-Shyn Chiang, Ron Davis, André Fialá, Barret Pfeiffer, Xi Rao, Carolina Rezaval, Iris Salecker for flies; DSHB (Iowa) for antibodies; AddGene for plasmids; Bloomington Stock Centre for *Drosophila* stocks. This work was funded by Marie-Curie Sklodowska Post-Doctoral fellowship to J.S.; Science Without Borders-CAPES PhD Studentship BEX 13380/13-3 to SUB; BBSRC Project Grants BB/R00871X/1 and BB/P004997/1 to A.H.; and Wellcome Trust Investigator Award 223197/Z/21/Z to A.H.

## AUTHOR CONTRIBUTIONS

J.S., V.C and A.H. designed and/or executed experiments and/or analysed data; S.U-B, G.L, D.S, X.W., F.R.C, M.M. executed experiments and analysed data; M.G.F, S.C., A.H. developed tools; D.K., G.J, V.C., A.H. supervised; J.S and A.H. wrote the manuscript. A.H. conceived and directed the project. All authors provided feedback on, and contributed to improvements to, the manuscript.

## DATA AVAILABILITY STATEMENT

All data are contained within the manuscript; metadata and statistical analysis details including full genotypes, sample sizes, statistical tests and p values have been provided in Table S2. This work generated fly stocks and molecular constructs, which we will distribute on request. Raw data will be distributed on request.

## MATERIALS AND METHODS

### Genetics

**Mutants:** *DNT2^37^* and *DNT2^18^*are protein null(Foldi et al. 2017, Ulian-Benitez et al. 2017). *Toll6^31^*is a null mutant allele(McIlroy et al. 2013). **Driver lines:** *DNT2-Gal4* is a CRISPR/Cas9-knock-in allele, with GAL4 at the start of the gene (this work, see below). *Toll-6-Gal4* was generated by RMCE from *MIMIC Toll-6^MIO2127^; kek6-Gal4* from *MIMIC Kek6^MI12953^*(Ulian-Benitez et al. 2017, Li et al. 2020). *MB320C-Gal4 (BSC68253), MB301B-Gal4 (BSC68311), R58E02-Gal4 (BSC41347), MB247-Gal80 (BSC64306), R58E02-LexA (BSC52740), R14C08-LexA (BSC52473)* were from the *Drosophila* Bloomington Stock Centre (BSC). *Dop1R2-LexA, Dop2R-LexA, Gad-LexA* were kindly provided by Yi Rao; *TH-LexA* (gift of Ron Davis), *TH-Gal4,R58E02-GAL4* (gift of Serge Birman); *G0431-Gal4 (DAL-GAL4), VT49239 nls LexA (DAL LexA)* (gifts from Ann-Shyn Chiang); tubGal80ts, Tdc-LexA. **Reporter lines:** *UAS-CD8::GFP,* for membrane tethered GFP; *UAS-histone-YFP*, for YFP-tagged nuclear histone; *UASFlybow1.1*, constant membrane tethered expression (gift of Iris Salecker); *13xLexAop-nls-tdTomato,* nuclear Tomato (gift of B. Pfeiffer); *UASCD8::RFP, LexAopCD8::GFP* (BSC32229), for dual binary expression; *UAS-homer-GCaMP* for post-synaptic densities (gift of André Fialá), *UAS-syt.eGFP* (BSC6925) and *LexAop-Syt-GCaMP* (BSC64413) for presynaptic sites. **For connectivity:** *UAS-DenMarkRFP, UAS-Dsyd1GFP* (gift of Carolina Rezaval); <u>TransTango:</u> *yw UAS-myrGFP,QUAS-mtdTomato-3xHA attP8; Trans-Tango@attP40* (BSC77124); <u>BAcTrace 806</u> *(w;LexAop2-Syb::GFP-P10(VK37) LexAop-QF2::SNAP25::HIVNES::Syntaxin(VK18)/CyO; UAS-B3Recombinase (attP2) UAS<B3Stop<BoNT/A (VK5) UAS<B3Stop<BoNT/A(VK27) QUAS-mtdTomato::HA/TM2*): **MCFO clones**: *hs-FLPG5.PEST;; 10xUAS (FRT.stop) myr::smGdP-OLLAS 10xUAS (FRT.stop) myr::smGdP-HA 10xUAS (FRT.stop) myr::smGdP-V5-THS-10xUAS (FRT.stop) myr::smGdP-FLAG* (BSC64086). **Optogenetic activation:** *20x UAS-V5-Syn-CsChrimson td tomato* (gift of B. Pfeiffer**). For thermogenetic activation:** *UASTrpA1@attP2 (BSC26264)* and *UAS-TrpA1@attP216 (BSC26263).* **UAS gene over-expression**: *UAS-DNT2CK, UAS-DNT2FL-GFP, UAS-DNT2-FL-47C, UAS-Toll-6^CY^*(McIlroy et al. 2013, Foldi et al. 2017). **UAS-RNAi knockdown:** *UAS-DNT2RNAi* (VDRC49195), *UAS-Toll6RNAi* (VDRC928), *y^1^v^1^; UAS-Toll6RNAi[Trip.HMS04251]* (BSC 56048*), UAS-kek6RNAi* (VDCR 109681), *y^1^v^1^; UAS-Dop2R-RNAi[TRiP.HMC02988](*BSC50621).

### Molecular biology

*DNT-2GAL4* was generated by CRISPR/Cas9 enhanced homologous recombination. 1 kb long 5’ and 3’ homology arms (HA) were amplified by PCR from genomic DNA of wild type flies, using primers for 5’HA: atcgcaccggtttttacaggcaccccatgtctga containing AgeI cutting site and cttgacgcggccgcTGTCAATTCATTCGCCGTCGAT containing NotI cutting site. 3’HA primers were: tattaggcgcgccATGACAAAAAGTATTAAACGTCCGCCC containing AscI cutting site and tactcgactagtgaagcacacccaaaatacccagg containing SpeI cutting site. HAs were sequentially cloned by conventional cloning into pGEM-T2AGal4 vector (Addgene #62894). For gRNA cloning, two 20 nucleotide gRNA oligos (gtcGACAAGTTCTTCTTACCTATG and aaacCATAGGTAAGAAGAACTTGTC) were designed using Optimal Target Finder. BbsI enzyme sites were added: gtc(g) at the 5’ end of sense oligo, and aaac at the 5’ end of the antisense oligo. gRNA is located at 41 bp downstream of the start codon of DNT-2 within the first coding exon. The gRNA was cloned into pU6.3 using conventional ligation. The two constructs were injected in Cas9 bearing flies and red fluorescent (3xP3-RFP) transformants were selected and balanced, after which 3xP3-RFP was removed with CRE-recombinase.

### qRT-PCR

qRT-PCR was carried out from 20 whole adult fruit-fly heads, frozen in liquid nitrogen before homogenising in 50 ml Trizol (Ambion #AM9738), followed by a stand RNA extraction protocol. RNA was treated with DNase treatment (ThermoFisher # AM1906) to remove genomic DNA. 200 ng of RNA was used for cDNA synthesis following Goscript Reverse Transcriptase (Promega #237815) protocol. Sample was diluted 1:4 with Nuclease free H_2_O. Standard qPCR was performed with SensiFast syb Green Mix (Bioline #B2092020) in ABI qPCR plate (GeneFlow #P3-0292) and machine. To amplify TH mRNA the following primers were used: TH-F: CGAGGACGAGATTTTGTTGGC and TH-R: TTGAGGCGGACCACCAAAG. GAPDH was used as a housekeeping control. Reactions were performed in triplicate. Specificity and size of amplification products were assessed by melting curve analyses. Target gene expression relative to reference gene is expressed as value of 2^-ΔΔCt^ (where Ct is the crossing threshold).

### Conditional expression

**Multiple Colour Flip Out clones:** *DNT2-Gal4, Toll6-Gal4* and *kek6-Gal4* were crossed with *hsFLP::PEST;;MCFO* flies, female offspring were collected and heat-shocked at 37°C in a water bath for 15 mins, then kept for 48h at 25°C before dissecting their brains. ***TransTango:*** DNT2-Gal4 or Oregon female virgins were crossed with *TransTango* males, progeny flies were raised at 18°C constantly and 15 days after eclosion, female flies were selected for immunostaining. **Thermogenetic activation with *TrpA1:*** Fruit-flies were bred at 18°C from egg laying to 4 days post-adult eclosion, then shifted to 29°C in a water bath for 24h followed by 24h recovery at room temperature for over-expressed *DNT-2FL-GFP*; for the other experiments, after breeding as above, adult flies were transferred to an incubator at 30°C, kept there for 24h and then brains were dissected. **Conditional gene over-expression and RNAi knockdown:** Flies bearing the temperature sensitive GAL4 repressor tubGal80^ts^ were kept at 18°C from egg laying to adult eclosion, then transferred to 30 °C incubator for 48h for Dcp-1+ and cell counting experiments and for 120h for TH+ cell counting.

### Immunostainings

Adult fruit-fly female brains were dissected (in PBS), fixed (in 4% para-formaldehyde, room temperature, 20-30min) and stained following standard protocols. Primary antibodies and their dilutions were as follows: Mouse anti-Brp (nc82) 1:10 (DSHB); Rabbit anti-GFP 1:250 (Thermofisher); Mouse anti-GFP 1:250 (Thermofisher); Chicken anti-GFP 1:500 (Aves); Rabbit anti-FLAG 1:50 (Sigma); Mouse anti-V5 1:50 (Invitrogen); Chicken anti-HA 1:50 (Aves); Rabbit anti-VGlut 1:500 (gift of Hermann); Mouse anti-TH 1:250 (Immunostar); Rabbit anti-TH 1:250 (Novus Biologicals); Rabbit anti-DsRed 1:250 (Clontek); Mouse anti-ChAT4B1 1:250 (DSHB); Rabbit anti-5-HT 1:500 (Immunostar); Rabbit Anti-DCP-1 1:250 (Cell Signalling). Seconday antibodies were all used at 1:500 and all were from Thermofisher: Alexa Flour 488 Goat anti-mouse, Alexa Flour 488 donkey anti-rabbit, Alexa Flour 488 Goat anti-rabbit (Fab’)2, Alexa Flour 488 Goat anti-chicken, Alexa Four 546 Goat anti-rabbit, Alexa Four 546 Goat anti-mouse, Alex Four 647 Goat anti-rabbit, Alex Four 647 Goat anti-mouse.

### Microscopy and Imaging

#### Laser scanning confocal microscopy

Stacks of microscopy images were acquired using laser scanning confocal microscopy with either Zeiss LSM710, 900 or Leica SP8. Brains were scanned with a resolution of 1024×1024, with Leica SP8 20x oil objective and 1 mm step for whole brain and DCP-1 stainings, 40x oil objective and 1 mm step for central brain. Resolution of 1024×512 was used for analysing PAM clusters with 0.96 mm step for cell counting; 0.5 mm step for neuronal morphology; and 63x oil objective with 0.5mm step for neuronal connections. Acquisition speed in Leica SP8 was 400Hz, with no line averaging. Resolution of 3072×3072 was used for single image analysis of synapses, using either Leica SP8 or Zeiss LSM900 and Airyscan acquisition with 40x water objective speed 6, and average 4, or with 1024×512, with 40x oil lens 2x zoom and 0.35mm step. TH counting in PAM were scanned with Zeiss 710 with a resolution 1024×1024, 40x oil objective, step 1 mm, speed 8. Zeiss LSM900 Airyscan with a resolution of 1024×1024, 40x water objective, speed 7, 0.7 zoom and 0.31µm step size was used for acquisition of optical sections of synapses in PAM neurons.

### Optogenetics and Epac1 FRET 2-photon imaging

To test whether DNT-2 neurons can respond to dopamine via the Dop2R inhibitory receptor, we used the cAMP sensor Epac1 and 2-photon confocal microscopy. Epac1 is FRET probe, whereby data are acquired from CFP and YFP emission and lower YFP/CFP ratio reveals higher cAMP levels. DNT-2Gal4 flies were crossed to *UAS-CsChrimson UAS Epac1* flies, to stimulate DNT-2 neurons and detect cAMP levels in DNT2 neurons. 1-3 day-old *DNT2Gal4>UASCsChrimson, UAS Epac1* flies were collected and separated in two groups. Flies bearing *DNT2Gal4 UASCsChrimson UAS Epac1 UASDop2RRNAi* were fed on 50μM all-trans retinal food for at least 3 days prior to imaging and kept in constant darkness prior to the experiment.

Optogenetic stimulation of fly brains expressing CsChrimson in DNT-2 neurons was carried out using a sapphire 588nm laser, in a 2-photon confocal microscope. For acquisition of YFP and CFP data from Epac1 samples, a FV30-FYC filter was applied, using a 925nm laser for both YFP and CFP imaging. The stimulation laser was targeted onto DNT-2 neuron projections in SMP region for 20s. Acquisition ROI was at DNT-2A cell bodies with a frame rate of around 10Hz. The first acquisition started 10s before the 20s stimulation and consequential acquisition was done every 30s for 10 cycles.

Image analysis of Epac data was carried out using ImageJ. The two channels (YFP and CFP) were separated, and the ratio of YFP/CFP for each pixel was calculated using the ImageJ>Image Calculator by diving YFP channel by CFP channel. The obtained result of YFP/CFP ratio was saved and the mean ratio of YFP/CFP in the ROI was calculated for each time point and 11 time points were used. The 11 values represent the ratio of YFP/CFP change in the cell body upon stimulation, with 30s interval and repeated 10 times.

#### Cell counting

To count cells labelled with nuclear reporters (e.g. Histone-YFP, nls-tdTomato) and Dcp-1+ cells, where signal is of high intensity, contrast and sphericity, we adapted the DeadEasy Central Brain ImageJ plug-in (Li et al. 2020) for automatic cell counting in adult brains. DeadEasy plug-ins automatically identify and count cells labelled with nuclear reporters in 3D stacks of confocal image in the nervous system of embryos (Forero et al. 2010a, Forero et al. 2010b), larvae (Kato et al. 2011, Forero et al. 2012, Losada-Perez et al. 2016, Harrison et al. 2021) and adult (Li et al. 2020) *Drosophila*. DeadEasy plug-ins are accurate at counting cells sparsely labelled with nuclear markers, and importantly, treat all genotypes objectively and equally yielding reliable data. Here, adult brains expressing Histone-YFP or nls-td-tomato reporters were dissected, fixed and scanned without staining them. DeadEasy Central Brain was used with threshold set to 75.

To count the TH labelled PAMs, where both the signal and the labelled-cell shape are more irregular, we used assisted manual cell counting using two methods. First, we developed a plug-in called DeadEasy DAN as follows. A median filter was used to reduce Poison noise, without having large losses at the edges. Then, a 3D morphological closing was performed. Next, all very dark pixels were assigned a value of zero. To mark each cell, each chasm in the image was found using a 3D extended h-minimal transform. As more than one local minimum can be found within each cell, which would result in counting a cell multiple times, a 3D inverse dome detection was performed, and then labelled. Thus, each inverse dome was used as a seed to identify each cell. Once the seeds were obtained, a 3D watershed transformation was performed to recover the shape of the cells. Then, we ran DeadEasy DAN on our raw data, to obtain a results stack of images and formed a merged stack between the raw and result stacks, to manually add any missing cells. This assisted cell counting method was effective at producing accurate cell counts with less labour and time than conventional manual counting and worked well for some genotypes (Figure 3D). However, it was less effective with RNAi knock-down genotypes, where the signal can be less intense, for which TH+ cells were counted manually, assisted by the ImageJ cell counter instead.

#### Dendrite analysis with Imaris

To analyze dendritic complexity, image data were processed with Imaris using the “Filaments” module with the default algorithm and “Autopatch function”. A simple region of interest (ROI) with parallelepiped shape was delimited. Thresholds were set for the largest and the smallest diameter of the dendrite, and this was consistent across samples within the same experiment. The starting point threshold was adjusted to only represent the soma of the neurons, and the “Seed Points threshold” to match the branches of the neurons. “Remove Seed Points Around Starting Points” and “Remove Disconnected Segments” were chosen, keeping the default values. The threshold for background substation and local contrast was consistent across all samples within an experiment. The “Edit function” within the Filaments module was used to correct any inaccuracy detected in the resulting tracing. Number of dendritic branches, dendritic segments, dendritic branch points and dendritic terminal points were collected to compare differences between the groups.

#### Vesicles, synapses and PSD analysis with Imaris

To analyse the number and volume of Homer-GCaMP GFP+ post-synaptic densities (PSD), Syt-GCaMP GFP+ presynaptic sites and DNT-2FLGFP+ vesicles, optical section images of confocal stacks through the brain were processed with the Imaris “Spot function”. To analyze the number and volume of Homer-GCaMP GFP+ post-synaptic densities (PSD), the “Surface module” from Imaris was used to restrict a region of interest (ROI). Then, “Absolute Intensity Thresholding” method was applied to each sample choosing the same cutoff each time. The resulting surface was applied to mask the original scan. The masked image was processed using “Image Processing module” from Imaris. Background subtraction followed by threshold cutoff filters were applied. Afterwards, the “Spots module” was used as explained below.

An ROI was determined for Syt-GCaMP GFP+ presynaptic sites and DNT-2FLGFP+ vesicles using the “Surface module” with the “Edit Manually” option “Algorithm”. The ROI for the SMP region started in the slide immediately after the last slide where the soma of PAM neurons was detectable, and finished in the last one where the dendrite was visible. The SMP region laterally was delimited by the black space given by the α lobe position. For the MB lobe region, the ROI started in the first slide where γ5,β’2 and β2 was visible (Aso et al., 2014) and finished in the last slide where this structure was appreciable. The surface was used to create a “Masked Channel”, which a posteriori was used to determine the spots using the Spots module.

The “Spots module” algorithm was set to “Different Spot Sizes”. An Estimated XY Diameter was set according to each experiment group, using the same within an experiment. “Background Subtraction” option was selected. “Intensity Center Filter” was used. “Spot Region” type was determined from “Local Contrast”, and the “Region Threshold” according to the “Region Border”. Setting of the threshold was consistent across genotypes.

### Behaviour

#### 1. Startle induced negative geotaxis Assay

was carried out as described in (Sun et al. 2018). Groups of approximately 10 male flies of the same genotype were placed in a fresh tube one night before the test, after which flies were transferred to a column formed with two empty tubes 15cm long and 2cm and then habituated for 30mins. Columns were tapped 3-4 times, flies fell to the bottom and then climbed upwards. Multiple rounds of testing were performed 3-7 times in a row per column. The process was filmed and films were analysed. Flies were scored during the first 15s after the tapping, and those that climbed above 13cm and those that climbed below 2cm were counted separately. Results given are mean ± SEM of the scores obtained with ten groups of flies per genotype. The performance index (PI) is defined as 1⁄2[(ntot + ntop − nbot)/ntot], where ntot, ntop, and nbot are the total number of flies, the number of flies at the top, and the number of flies at the bottom, respectively. The assay was carried out at 25°C, 55% humidity. Flies with tubGal80^ts^ to conditionally overexpress or knock-down were shifted from 18 to 30°at eclosion and kept for 5 days at 30°C to induce Gal4. Experiments were carried in an environmental chamber at 31°C, 60% humidity or in a humidity and temperature-controlled behaviour lab always kept at 25°C.

#### 2. Spontaneous Locomotion in an open arena

Male flies of each genotype were collected and kept in groups of 10-20 flies, in vials containing fresh fly food for 5-9 days. Before filming, three male flies from one genotype were transferred into a 24 mm well of a multi-well plate using an aspirator and habituated for 15 – 20 mins. The multi-well plate with transparent lid and bottom was placed on a white LED light pad (XIAOSTAR Light Box) and either inside a light-shielding black box (PULUZ, 40*40*40cm) in a room with constant temperature (25°C) and humidity (55%) to maintain stable environmental conditions (Figure 7D), or inside a temperature controlled environmental chamber at 18°C or 30°C (Figure 7E,F,). The locomotion behaviour of freely-moving flies was filmed with a camera (Panasonic, HC-V260) in the morning from ZT1-ZT4 and for 10 min at a frame-rate of 25 fps. The 10-min videos were trimmed (from 00:02:00 to 00:07:00) to 5-min videos for analysis using Flytracker software(Eyjolfsdottir et al. 2014). For the *DNT-2* mutants and over-expression of Toll-6^CY^ experiments, flies were bred and tested at 25 °C. To test over-expression of *DNT-2* with *tubGAL80^ts^; DNT2-Gal4>UAS-DNT-2FL*, flies were raised at 18°C until eclosion, and controls were kept and tested at 18°C; test groups were transferred directly after eclosion to 30°C for 5 days and were tested in an environmental chamber kept at 30°C, 60%. For thermo-genetic activation of DNT2 neurons using TrpA1 (DNT-2GAL4>UASTrpA1), flies were bred at 18°C and kept at 18°C for 7-9 days post-eclosion. Following habituation at 18°c for 20 mins in the multi-well plates, they were transferred to the 30°C chamber 10 mins before filming to activate TrpA1 and then filmed for the following 10 minutes. Fly locomotion activity was tracked using FlyTracker and calculated (distance and speed) in Matlab(Eyjolfsdottir et al. 2014) using the raw data generated from the tracking procedure. The “walking distance” was calculated as the sum of the distance flies moved and the “walking speed” was the speed of flies only when they were walking, and it was calculated using only frames where flies moved above 4 mm/s (which corresponds to two body lengths).

#### 3. Appetitive long term memory test

Appetitive long-term memory was tested as described in Krashes and Waddell(Krashes and Waddell 2011). The two conditioning odours used were isoamyl acetate, Sigma-Aldrich #24900822, 6mL in 8mL mineral oil (Sigma-Aldrich #330760) and 4-methylcyclohexanol (Sigma-Aldrich # 153095, 10mL in 8mL mineral oil). Groups of 80-120 mixed sex flies were starved in a 1% agar tube filled with a damp 20 x 60 mm piece of filter paper for 18-20 hour before conditioning. During conditioning training, one odorant was presented with a dry filter paper (unconditioned odour, CS-) for 2 minutes, before a 30 second break, and presentation of a second odorant with filter paper coated with dry sucrose (conditioned odour, CS+). The test was repeated pairing the other odorant with sucrose, with a different group of flies to form one replicate. After training, flies were transferred back to agar tubes for testing 24 hours later. Performance index (PI) was calculated in the same way as in Krashes and Waddell(Krashes and Waddell 2011), as the number of flies approaching the conditioned odour minus the number of flies going in the opposite direction, divided by the total number of flies. A single PI values is the average score from the test with the reverse conditioning odour combination. Groups for which the total number of flies among both odorants was below 15 were discarded. For the *DNT-2* over-expression experiments with *tubGAL80^ts^; DNT-2>DNT-2FL,* flies were raised at 18°C until 7-9 days post-eclosion. They were then either transferred to and maintained at 23°C (controls) or 30°C for 18-20h starvation, training, and up to testing 24h later.

### Statistical analysis

Statistical analyses were carried out using Graph-Pad Prism. Confidence interval was 95%, setting significance at p<0.05. Chi-square tests were carried out when comparing categorical data. Numerical data were tested first for their type of distributions. If data were distributed normally, unpaired Student t-tests were used to compare means between two groups and One Way ANOVA or Welch ANOVA for larger groups, followed by post-doc Dunnett test for multiple comparisons to a fixed control. Two Way ANOVA was used when comparisons to two variables were made. If data were not normally distributed, non-parametric Mann-Whitney U-test for 2 two group comparisons and Kruskal Wallis ANOVA for larger groups, followed by post-hoc Dunn’s multiple comparisons test to a fixed control. Statistical details including full genotypes, sample sizes, tests and p values are provided in Supplementary Table S2.

## SUPPLEMENTAL INFORMATION

**Supplementary Figure S1 Identification of neurotransmitter type for DNT-2A neurons. (A)** All four DNT-2A neurons per hemibrain are glutamatergic as DNT-2>FlyBow1.1 reporter co-localises with anti-vGlut in these neurons as well as in their projections at SMP (arrows point at cluster of 4 cells). **(B)** All four DNT-2A neurons per hemibrain have Dop2R (arrows point to cluster, each neuron seen in magenta). **(C, D)** Anterior DNT-2A neurons projecting at SMP are not dopaminergic, as there was no co-localisation between *DNT-2>histoneYFP* and anti-TH in the anterior brain **(C),** and there was no colocalization in other DNT-2+ neurons in the posterior brain either **(D).** Higher magnification of dotted boxes on the right. **(E)** There was no colocalisation with *Dop1R2LexA>CD8::GFP, DNT-2Gal4>CD8::RFP* either. **(F)** DNT-2 neurons are not serotonergic, as there was no co-localization between *DNT-2>FlyBow1.1* and the serotoninergic neuron marker anti-5HT**. (G)** They are not octopaminergic, as there was no overlap between *TdcLexA>mCD8::GFP and DNT-2Gal4>CD8::RFP*. **(H)** There was no overlap between *DNT-2Gal4>histoneYFP* and the cholinergic neuron marker anti-ChAT4b1. **(I,J)** Anterior DNT-2A neurons are not GABAergic, but lateral DNT-2 neurons are, as visualised with *DNT-2>CD8-RFP, GADLexA>CD8-GFP*. Scale bars: (A right,B, F, G, H, I, J) 20µm; (C,D) 50 µm. For further genotypes and sample sizes, see Supplementary Table S2.

**Supplementary Figure S2 Identification of *Toll-6* and *kek-6* expressing neurons. (A)** *Toll-6Gal4, MB247-Gal80>UASFlyBow1.1* and *kek-6Gal4, MB247-Gal80>UASFlyBow1.1* revealed Toll-6+ and Kek-6+ projections on MB g2a’1. **(B)** *R25D01LexA>CD8GFP, kek6-Gal4>CD8RFP* overlapped in MB g2a’1. **(C)** *R14C08LexA>CD8GFP, Toll-6Gal4>CD8-RFP13* revealed co-localization in MBON-M4/M6 cell bodies. **(D,E)** Co-localisation of *Toll-6>Histone-YFP* and *kek-6>HistoneYFP* with TH in PPL1 neurons. **(F)** Number of PAM and PPL1 neurons expressing *Tolls* and *kek-6,* revealed from published RNAseq data (Scope, see also Table S1). **(G, H)** Toll-6 and kek-6 are expressed in DAL neurons. **(G)** DAL neurons express *Toll-6,* as revealed by co-localization between *DAL-LexA (VT49239-LexA)>mCD8-GFP* and *Toll-6GAL4>mCD8-RFP*. **(H)** DAL neurons express *kek-6*, as revealed by colocalization between *DAL-LexA (VT49239-LexA)>mCD8-GFP* and *kek6GAL4>mCD8RFP*. **(I, J)** Toll-6 and kek-6 are expressed in MB neurons. **(I)** *Toll-6>MCFO* clones reveal expression at least in MB neurons g medial (gm), abcore (abc) and absurface (abs). **(K)** *kek-6>MCFO* clones reveal expression at least in MB Kenyon cells aba’b’g formed and MB g2a’1 and a2a’2 and PPL1-γ2α’1 and PPL1-α2α’2. Scale bars: (B, H) 50µm; (A,C,D,G, I, J) 30µm. For further genotypes and sample sizes, see Supplementary Table S2.

**Supplementary Figure S3 TransTango connectivity controls. (A)** *UAS-TransTango/+* control, showing background GFP expression in mushroom bodies and Tomato+ signal in sub-aesophageal ganglion (SOG). **(B)** *DNT-2GAL4>UAS-TransTango*, revealing the GFP+ DNT-2 neurons and Tomato+ signal in DNT-2 neurons. TransTango also reveals what appears to be feedback connections between DNT-2 neurons from SMP to PRW (arrow). These are controls for Figure 2A. Scale bars: (A,B) 50µm. For further genotypes and sample sizes, see Supplementary Table S2.

**Supplementary Figure S4 Neuronal activity increased production and cleavage of DNT-2. (A)** Thermogenetic activation of DNT-2 neurons increased the number of DNT-2FLGFP+ vesicles or spots (*DNT-2>DNT-2-FL-GFP*, *TrpA1* with anti-GFP). Control 18°C: Unpaired Student t test, ns. Activation 30°C 24h: Mann Whitney**. (B)** Western blot from fly heads over-expressing DNT-2FL-GFP in DNT-2 neurons showing that neuronal activation with TrpA1 at 30°C increased DNT-2FL-GFP cleavage. (Genotype: *DNT-2>TrpA1, DNT-2FL-GFP*). Controls are flies of the same genotype kept constantly at 18°C as well as flies treated also at 30°C but lacking TrpA1. High temperature (30°C) is sufficient to increase fly activity. ***p<0.001. ns= not significant. For further genotypes, sample sizes, p values and other statistical details see Supplementary Table S2.

**Supplementary Figure S5 Controls for Figure 5B**. Control at 18°C, for Figure 5B. DNT-2 bidirectional arborisation at SMP was visualised with Homer-GCaMP and anti-GFP antibodies (*DNT-21>homer-GCAMP* and *DNT-2>homer-GCAMP, TrpA1* at 18°C). Unpaired Student t-test, ns. Scale bars: (A,B) 30µm. For further genotypes, sample sizes, p values and other statistical details see Supplementary Table S2.

**Supplementary Figure S6 Locomotion in open arena controls. (A)** Controls for Figure 6E. In 18°C controls, GAL80 is on and GAL4 is off, thus there is no adult specific *DNT-2FL* overexpression in DNT-2 neurons in these flies. (Genotype: *tubGAL80ts, DNT-2>DNT-2FL*). Consistently, there was no increase in locomotion speed in an open arena, in flies over-expressing *DNT-2FL* relative to controls, compare also to Figure 6E. Kruskal Wallis ANOVA ns (lower median in *UAS-DNT-2FL/+* control). **(B)** Controls for Figure 6F. The cation TrpA1 opens at high temperatures (e.g. 30°C) but remains closed at 18°C. Consistently, there was no effect in locomotion at 18°C in flies of genotype *DNT-2>TrpA1* compared to controls, compare also to 30°C data in Figure 6F. Kruskal Wallis ANOVA ns. For further genotypes, sample sizes, p values and other statistical details see Supplementary Table S2.

**Supplementary Figure S7 Altering *DNT-2* levels induced seizures.** Knock-down or over-expression of *DNT-2* in the adult, using *GAL80^ts^*. Fruit-flies were reared at 18°C from egg laying to adult eclosion, when they were transferred to 30°C and kept there for 5 days prior to testing. To test for seizures, we used the bang-sensitivity test. *DNT2^37^/DNT2^18^*mutant flies (left) and adult *DNT-2* knock-down flies (*tubGal80^ts^ DNT-2Gal4>DNT-2RNAi,* centre) took longer to recover than controls. Over-expression of *DNT-2FL* in DNT-2 neurons (*tubGal80^ts^ DNT-2Gal4>DNT-2FL,* right) increased variability in recovery time. For further genotypes and sample sizes see Supplementary Table S2.

**Table S1 Expression of *Tolls, keks* and Toll downstream adaptors in cells related to DNT-2A neurons.** Genes expressed in DNT-2 neurons, their potential and/or experimentally verified inputs and outputs, were identified with a combination of reporters (this work) and data from public single-cell RNAseq databases.

**Table S2 Statistical analysis.** Table providing full genotypes, sample sizes, statistical tests, multiple comparison corrections and p values.

## REFERENCES

Adel, M. and L. C. Griffith (2021). “The Role of Dopamine in Associative Learning in Drosophila: An Updated Unified Model.” Neurosci Bull 37(6): 831–852.

Adhikarla, S. V., N. K. Jha, V. K. Goswami, A. Sharma, A. Bhardwaj, A. Dey, C. Villa, Y. Kumar and S. K. Jha (2021). “TLR-Mediated Signal Transduction and Neurodegenerative Disorders.” Brain Sci 11(11).

Anthoney, N., I. Foldi and A. Hidalgo (2018). “Toll and Toll-like receptor signalling in development.” Development 145(9).

Arikkath, J. (2012). “Molecular mechanisms of dendrite morphogenesis.” Front Cell Neurosci 6: 61.

Arnot, C. J., N. J. Gay and M. Gangloff (2010). “Molecular mechanism that induces activation of Spatzle, the ligand for the Drosophila Toll receptor.” J Biol Chem 285(25): 19502–19509.

Aso, Y., D. Hattori, Y. Yu, R. M. Johnston, N. A. Iyer, T. T. Ngo, H. Dionne, L. F. Abbott, R. Axel, H. Tanimoto and G. M. Rubin (2014a). “The neuronal architecture of the mushroom body provides a logic for associative learning.” Elife 3: e04577.

Aso, Y., A. Herb, M. Ogueta, I. Siwanowicz, T. Templier, A. B. Friedrich, K. Ito, H. Scholz and H. Tanimoto (2012). “Three dopamine pathways induce aversive odor memories with different stability.” PLoS Genet 8(7): e1002768.

Aso, Y., D. Sitaraman, T. Ichinose, K. R. Kaun, K. Vogt, G. Belliart-Guerin, P. Y. Placais, A. A. Robie, N. Yamagata, C. Schnaitmann, W. J. Rowell, R. M. Johnston, T. T. Ngo, N. Chen, W. Korff, M. N. Nitabach, U. Heberlein, T. Preat, K. M. Branson, H. Tanimoto and G. M. Rubin (2014b). “Mushroom body output neurons encode valence and guide memory-based action selection in Drosophila.” Elife 3: e04580.

Baltruschat, L., L. Prisco, P. Ranft, J. S. Lauritzen, A. Fiala, D. D. Bock and G. Tavosanis (2021). “Circuit reorganization in the Drosophila mushroom body calyx accompanies memory consolidation.” Cell Rep 34(11): 108871.

Barth, M. and M. Heisenberg (1997). “Vision affects mushroom bodies and central complex in Drosophila melanogaster.” Learn Mem 4(2): 219–229.

Barth, M., H. V. Hirsch, I. A. Meinertzhagen and M. Heisenberg (1997). “Experience-dependent developmental plasticity in the optic lobe of Drosophila melanogaster.” J Neurosci 17(4): 1493–1504.

Berton, O., C. A. McClung, R. J. Dileone, V. Krishnan, W. Renthal, S. J. Russo, D. Graham, N. M. Tsankova, C. A. Bolanos, M. Rios, L. M. Monteggia, D. W. Self and E. J. Nestler (2006). “Essential role of BDNF in the mesolimbic dopamine pathway in social defeat stress.” Science 311(5762): 864–868.

Bharmauria, V., A. Ouelhazi, R. Lussiez and S. Molotchnikoff (2022). “Adaptation-induced plasticity in the sensory cortex.” J Neurophysiol 128(4): 946–962.

Blochl, A. and C. Sirrenberg (1996). “Neurotrophins stimulate the release of dopamine from rat mesencephalic neurons via Trk and p75Lntr receptors.” J Biol Chem 271(35): 21100–21107.

Bolus, H., K. Crocker, G. Boekhoff-Falk and S. Chtarbanova (2020). “Modeling Neurodegenerative Disorders in Drosophila melanogaster.” Int J Mol Sci 21(9).

Boto, T., T. Louis, K. Jindachomthong, K. Jalink and S. M. Tomchik (2014). “Dopaminergic modulation of cAMP drives nonlinear plasticity across the Drosophila mushroom body lobes.” Curr Biol 24(8): 822–831.

Boto, T., A. Stahl and S. M. Tomchik (2020). “Cellular and circuit mechanisms of olfactory associative learning in Drosophila.” J Neurogenet 34(1): 36–46.

Bushey, D., G. Tononi and C. Cirelli (2011). “Sleep and synaptic homeostasis: structural evidence in Drosophila.” Science 332(6037): 1576–1581.

Cachero, S., M. Gkantia, A. S. Bates, S. Frechter, L. Blackie, A. McCarthy, B. Sutcliffe, A. Strano, Y. Aso and G. S. X. E. Jefferis (2020). “BAcTrace a new tool for retrograde tracing of neuronal circuit.” Nature Methods 17: 1254–1261.

Carim-Todd, L., K. G. Bath, G. Fulgenzi, S. Yanpallewar, D. Jing, C. A. Barrick, J. Becker, H. Buckley, S. G. Dorsey, F. S. Lee and L. Tessarollo (2009). “Endogenous truncated TrkB.T1 receptor regulates neuronal complexity and TrkB kinase receptor function in vivo.” J Neurosci 29(3): 678–685.

Casarotto, P. C., M. Girych, S. M. Fred, V. Kovaleva, R. Moliner, G. Enkavi, C. Biojone, C. Cannarozzo, M. P. Sahu, K. Kaurinkoski, C. A. Brunello, A. Steinzeig, F. Winkel, S. Patil, S. Vestring, T. Serchov, C. Diniz, L. Laukkanen, I. Cardon, H. Antila, T. Rog, T. P. Piepponen, C. R. Bramham, C. Normann, S. E. Lauri, M. Saarma, I. Vattulainen and E. Castren (2021). “Antidepressant drugs act by directly binding to TRKB neurotrophin receptors.” Cell 184(5): 1299–1313 e1219.

Castren, E. and L. Monteggia (2021). “Brain-Derived Neurotrophic Factor Signaling in Depression and Antidepressant Action.” Biol Psychiatry.

Chan, R. F., G. Turecki, A. A. Shabalin, J. Guintivano, M. Zhao, L. Y. Xie, G. van Grootheest, Z. A. Kaminsky, B. Dean, B. Penninx, K. A. Aberg and E. van den Oord (2020). “Cell Type-Specific Methylome-wide Association Studies Implicate Neurotrophin and Innate Immune Signaling in Major Depressive Disorder.” Biol Psychiatry 87(5): 431–442.

Chen, C. C. and J. C. Brumberg (2021). “Sensory Experience as a Regulator of Structural Plasticity in the Developing Whisker-to-Barrel System.” Frontiers in Cellular Neuroscience 15.

Chen, C. C., J. K. Wu, H. W. Lin, T. P. Pai, T. F. Fu, C. L. Wu, T. Tully and A. S. Chiang (2012). “Visualizing long-term memory formation in two neurons of the Drosophila brain.” Science 335(6069): 678–685.

Chen, C. Y., Y. C. Shih, Y. F. Hung and Y. P. Hsueh (2019). “Beyond defense: regulation of neuronal morphogenesis and brain functions via Toll-like receptors.” J Biomed Sci 26(1): 90.

Cheng, L. K., Y. H. Chiu, Y. C. Lin, W. C. Li, T. Y. Hong, C. J. Yang, C. H. Shih, T. C. Yeh, W. I. Tseng, H. Y. Yu, J. C. Hsieh and L. F. Chen (2022). “Long-term musical training induces white matter plasticity in emotion and language networks.” Hum Brain Mapp.

Coban, B., H. Poppinga, E. Y. Rachad, B. Geurten, D. Vasmer, F. J. Rodriguez Jimenez, Y. Gadgil, S. H. Deimel, I. Alyagor, O. Schuldiner, I. C. Grunwald Kadow, T. D. Riemensperger, A. Widmann and A. Fiala (2024). “The caloric value of food intake structurally adjusts a neuronal mushroom body circuit mediating olfactory learning in Drosophila.” Learn Mem 31(5).

Costa, M., J. D. Manton, A. D. Ostrovsky, S. Prohaska and G. S. Jefferis (2016). “NBLAST: Rapid, Sensitive Comparison of Neuronal Structure and Construction of Neuron Family Databases.” Neuron 91(2): 293–311.

Cotman, C. W. and N. C. Berchtold (2002). “Exercise: a behavioral intervention to enhance brain health and plasticity.” Trends Neurosci 25(6): 295–301.

Croset, V., C. D. Treiber and S. Waddell (2018). “Cellular diversity in the Drosophila midbrain revealed by single-cell transcriptomics.” Elife 7.

DeLotto, Y. and R. DeLotto (1998). “Proteolytic processing of the Drosophila Spatzle protein by easter generates a dimeric NGF-like molecule with ventralising activity.” Mech Dev 72(1-2): 141–148.

Devaud, J. M., A. Acebes and A. Ferrus (2001). “Odor exposure causes central adaptation and morphological changes in selected olfactory glomeruli in Drosophila.” J Neurosci 21(16): 6274–6282.

Duhart, J. M., A. Herrero, G. de la Cruz, J. I. Ispizua, N. Pirez and M. F. Ceriani (2020). “Circadian Structural Plasticity Drives Remodeling of E Cell Output.” Curr Biol 30(24): 5040–5048 e5045.

Ernst, C., V. Deleva, X. Deng, A. Sequeira, A. Pomarenski, T. Klempan, N. Ernst, R. Quirion, A. Gratton, M. Szyf and G. Turecki (2009). “Alternative splicing, methylation state, and expression profile of tropomyosin-related kinase B in the frontal cortex of suicide completers.” Arch Gen Psychiatry 66(1): 22–32.

Eyjolfsdottir, E., S. Branson, X. P. Burgos-Artizzu, E. D. Hoopfer, J. Schor, D. J. Anderson and P. Perona (2014). Detecting Social Actions of Fruit Flies, Cham, Springer International Publishing.

Feany, M. B. and W. W. Bender (2000). “A Drosophila model of Parkinson’s disease.” Nature 404(6776): 394–398.

Feldman, D. E. and M. Brecht (2005). “Map plasticity in somatosensory cortex.” Science 310(5749): 810–815.

Fenner, B. M. (2012). “Truncated TrkB: beyond a dominant negative receptor.” Cytokine Growth Factor Rev 23(1-2): 15–24.

Fernandez, M. P., J. Berni and M. F. Ceriani (2008). “Circadian remodeling of neuronal circuits involved in rhythmic behavior.” PLoS Biol 6(3): e69.

Ferrer, I., C. Marin, M. J. Rey, T. Ribalta, E. Goutan, R. Blanco, E. Tolosa and E. Marti (1999). “BDNF and full-length and truncated TrkB expression in Alzheimer disease. Implications in therapeutic strategies.” J Neuropathol Exp Neurol 58(7): 729–739.

Figueroa-Hall, L. K., M. P. Paulus and J. Savitz (2020). “Toll-Like Receptor Signaling in Depression.” Psychoneuroendocrinology 121: 104843.

Foldi, I., N. Anthoney, N. Harrison, M. Gangloff, B. Verstak, M. P. Nallasivan, S. AlAhmed, B. Zhu, M. Phizacklea, M. Losada-Perez, M. Moreira, N. J. Gay and A. Hidalgo (2017). “Three-tier regulation of cell number plasticity by neurotrophins and Tolls in Drosophila.” J Cell Biol 216(5): 1421–1438.

Forero, M. G., K. Kato and A. Hidalgo (2012). “Automatic cell counting in vivo in the larval nervous system of Drosophila.” J Microsc 246(2): 202–212.

Forero, M. G., A. R. Learte, S. Cartwright and A. Hidalgo (2010a). “DeadEasy Mito-Glia: automatic counting of mitotic cells and glial cells in Drosophila.” PLoS One 5(5): e10557.

Forero, M. G., J. A. Pennack and A. Hidalgo (2010b). “DeadEasy neurons: automatic counting of HB9 neuronal nuclei in Drosophila.” Cytometry A 77(4): 371–378.

Forero, M. G., J. A. Pennack, A. R. Learte and A. Hidalgo (2009). “DeadEasy caspase: automatic counting of apoptotic cells in Drosophila.” PLoS One 4(5): e5441.

Forrest, M. P., E. Parnell and P. Penzes (2018). “Dendritic structural plasticity and neuropsychiatric disease.” Nat Rev Neurosci 19(4): 215–234.

Fryer, R. H., D. R. Kaplan, S. C. Feinstein, M. J. Radeke, D. R. Grayson and L. F. Kromer (1996). “Developmental and mature expression of full-length and truncated TrkB receptors in the rat forebrain.” J Comp Neurol 374(1): 21–40.

Fuentes-Medel, Y., M. A. Logan, J. Ashley, B. Ataman, V. Budnik and M. R. Freeman (2009). “Glia and muscle sculpt neuromuscular arbors by engulfing destabilized synaptic boutons and shed presynaptic debris.” PLoS Biol 7(8): e1000184.

Gage, F. H. (2019). “Adult neurogenesis in mammals.” Science 364(6443): 827–828.

Garrett, M. E., X. J. Qin, D. Mehta, M. F. Dennis, C. E. Marx, G. A. Grant, V. A. M.-A. M. Workgroup, P. Initiative, Injury, C. Traumatic Stress Clinical, P. G. Psychiatric Genomics Consortium, M. B. Stein, N. A. Kimbrel, J. C. Beckham, M. A. Hauser and A. E. Ashley-Koch (2021). “Gene Expression Analysis in Three Posttraumatic Stress Disorder Cohorts Implicates Inflammation and Innate Immunity Pathways and Uncovers Shared Genetic Risk With Major Depressive Disorder.” Front Neurosci 15: 678548.

Gay, N. J. and M. Gangloff (2007). “Structure and function of Toll receptors and their ligands.” Annu Rev Biochem 76: 141–165.

Gorska-Andrzejak, J., A. Keller, T. Raabe, L. Kilianek and E. Pyza (2005). “Structural daily rhythms in GFP-labelled neurons in the visual system of Drosophila melanogaster.” Photochem Photobiol Sci 4(9): 721–726.

Guillin, O., J. Diaz, P. Carroll, N. Griffon, J. C. Schwartz and P. Sokoloff (2001). “BDNF controls dopamine D3 receptor expression and triggers behavioural sensitization.” Nature 411(6833): 86–89.

Guven-Ozkan, T. and R. L. Davis (2014). “Functional neuroanatomy of Drosophila olfactory memory formation.” Learn Mem 21(10): 519–526.

Harrison, N. J., E. Connolly, A. Gascon Gubieda, Z. Yang, B. Altenhein, M. Losada Perez, M. Moreira, J. Sun and A. Hidalgo (2021). “Regenerative neurogenic response from glia requires insulin-driven neuron-glia communication.” Elife 10.

Hearn, M. G., Y. Ren, E. W. McBride, I. Reveillaud, M. Beinborn and A. S. Kopin (2002). “A Drosophila dopamine 2-like receptor: Molecular characterization and identification of multiple alternatively spliced variants.” Proc Natl Acad Sci U S A 99(22): 14554–14559.

Heisenberg, M. (2003). “Mushroom body memoir: from maps to models.” Nat Rev Neurosci 4(4): 266–275.

Heisenberg, M., M. Heusipp and C. Wanke (1995). “Structural plasticity in the Drosophila brain.” J Neurosci 15(3 Pt 1): 1951–1960.

Hepburn, L., T. K. Prajsnar, C. Klapholz, P. Moreno, C. A. Loynes, N. V. Ogryzko, K. Brown, M. Schiebler, K. Hegyi, R. Antrobus, K. L. Hammond, J. Connolly, B. Ochoa, C. Bryant, M. Otto, B. Surewaard, S. L. Seneviratne, D. M. Grogono, J. Cachat, T. Ny, A. Kaser, M. E. Torok, S. J. Peacock, M. Holden, T. Blundell, L. Wang, P. Ligoxygakis, L. Minichiello, C. G. Woods, S. J. Foster, S. A. Renshaw and R. A. Floto (2014). “Innate immunity. A Spaetzle-like role for nerve growth factor beta in vertebrate immunity to Staphylococcus aureus.” Science 346(6209): 641–646.

Hoffmann, A., A. Funkner, P. Neumann, S. Juhnke, M. Walther, A. Schierhorn, U. Weininger, J. Balbach, G. Reuter and M. T. Stubbs (2008a). “Biophysical characterization of refolded Drosophila Spatzle, a cystine knot protein, reveals distinct properties of three isoforms.” J Biol Chem 283(47): 32598–32609.

Hoffmann, A., P. Neumann, A. Schierhorn and M. T. Stubbs (2008b). “Crystallization of Spatzle, a cystine-knot protein involved in embryonic development and innate immunity in Drosophila melanogaster.” Acta Crystallogr Sect F Struct Biol Cryst Commun 64(Pt 8): 707–710.

Holtmaat, A. and K. Svoboda (2009). “Experience-dependent structural synaptic plasticity in the mammalian brain.” Nat Rev Neurosci 10(9): 647–658.

Horch, H. W. and L. C. Katz (2002). “BDNF release from single cells elicits local dendritic growth in nearby neurons.” Nat Neurosci 5(11): 1177–1184.

Hu, X., Y. Yagi, T. Tanji, S. Zhou and Y. T. Ip (2004). “Multimerization and interaction of Toll and Spatzle in Drosophila.” Proc Natl Acad Sci U S A 101(25): 9369–9374.

Huang, C., J. Luo, S. J. Woo, L. A. Roitman, J. Li, V. A. Pieribone, M. Kannan, G. Vasan and M. J. Schnitzer (2024). “Dopamine-mediated interactions between short- and long-term memory dynamics.” Nature.

Hyman, C., M. Hofer, Y. A. Barde, M. Juhasz, G. D. Yancopoulos, S. P. Squinto and R. M. Lindsay (1991). “BDNF is a neurotrophic factor for dopaminergic neurons of the substantia nigra.” Nature 350(6315): 230–232.

Jenks, K. R., K. Tsimring, J. P. K. Ip, J. C. Zepeda and M. Sur (2021). “Heterosynaptic Plasticity and the Experience-Dependent Refinement of Developing Neuronal Circuits.” Front Neural Circuits 15: 803401.

Kato, K., M. G. Forero, J. C. Fenton and A. Hidalgo (2011). “The glial regenerative response to central nervous system injury is enabled by pros-notch and pros-NFkappaB feedback.” PLoS Biol 9(8): e1001133.

Krashes, M. J. and S. Waddell (2008). “Rapid consolidation to a radish and protein synthesis-dependent long-term memory after single-session appetitive olfactory conditioning in Drosophila.” J Neurosci 28(12): 3103–3113.

Krashes, M. J. and S. Waddell (2011). “Drosophila appetitive olfactory conditioning.” Cold Spring Harb Protoc 2011(5): pdb prot5609.

Kremer, M. C., F. Christiansen, F. Leiss, M. Paehler, S. Knapek, T. F. Andlauer, F. Forstner, P. Kloppenburg, S. J. Sigrist and G. Tavosanis (2010). “Structural long-term changes at mushroom body input synapses.” Curr Biol 20(21): 1938–1944.

Krishnan, V. and E. J. Nestler (2008). “The molecular neurobiology of depression.” Nature 455(7215): 894–902.

Kuner, R. and H. Flor (2016). “Structural plasticity and reorganisation in chronic pain.” Nat Rev Neurosci 18(1): 20–30.

Lee, J. L., B. J. Everitt and K. L. Thomas (2004). “Independent cellular processes for hippocampal memory consolidation and reconsolidation.” Science 304(5672): 839–843.

Leemhuis, E., L. De Gennaro and A. M. Pazzaglia (2019). “Disconnected Body Representation: Neuroplasticity Following Spinal Cord Injury.” J Clin Med 8(12).

Levi-Montalcini, R., S. D. Skaper, R. Dal Toso, L. Petrelli and A. Leon (1996). “Nerve growth factor: from neurotrophin to neurokine.” Trends Neurosci 19(11): 514–520.

Lewis, M., C. J. Arnot, H. Beeston, A. McCoy, A. E. Ashcroft, N. J. Gay and M. Gangloff (2013). “Cytokine Spatzle binds to the Drosophila immunoreceptor Toll with a neurotrophin-like specificity and couples receptor activation.” Proc Natl Acad Sci U S A 110(51): 20461–20466.

Li, G., M. G. Forero, J. S. Wentzell, I. Durmus, R. Wolf, N. C. Anthoney, M. Parker, R. Jiang, J. Hasenauer, N. J. Strausfeld, M. Heisenberg and A. Hidalgo (2020). “A Toll-receptor map underlies structural brain plasticity.” Elife 9.

Li, G. and A. Hidalgo (2021). “The Toll Route to Structural Brain Plasticity.” Front Physiol 12: 679766.

Linneweber, G. A., M. Andriatsilavo, S. B. Dutta, M. Bengochea, L. Hellbruegge, G. Liu, R. K. Ejsmont, A. D. Straw, M. Wernet, P. R. Hiesinger and B. A. Hassan (2020). “A neurodevelopmental origin of behavioral individuality in the Drosophila visual system.” Science 367(6482): 1112–1119.

Liu, C., P. Y. Placais, N. Yamagata, B. D. Pfeiffer, Y. Aso, A. B. Friedrich, I. Siwanowicz, G. M. Rubin, T. Preat and H. Tanimoto (2012). “A subset of dopamine neurons signals reward for odour memory in Drosophila.” Nature 488(7412): 512–516.

Losada-Perez, M., N. Harrison and A. Hidalgo (2016). “Molecular mechanism of central nervous system repair by the Drosophila NG2 homologue kon-tiki.” J Cell Biol 214(5): 587–601.

Lu, B., G. Nagappan, X. Guan, P. J. Nathan and P. Wren (2013). “BDNF-based synaptic repair as a disease-modifying strategy for neurodegenerative diseases.” Nat Rev Neurosci 14(6): 401–416.

Lu, B., P. T. Pang and N. H. Woo (2005). “The yin and yang of neurotrophin action.” Nat Rev Neurosci 6(8): 603–614.

Ma, Y., J. Li, I. Chiu, Y. Wang, J. A. Sloane, J. Lu, B. Kosaras, R. L. Sidman, J. J. Volpe and T. Vartanian (2006). “Toll-like receptor 8 functions as a negative regulator of neurite outgrowth and inducer of neuronal apoptosis.” J Cell Biol 175(2): 209–215.

MacLaren, C. M., T. A. Evans, D. Alvarado and J. B. Duffy (2004). “Comparative analysis of the Kekkon molecules, related members of the LIG superfamily.” Dev Genes Evol 214(7): 360–366.

Maguire, E. A., D. G. Gadian, I. S. Johnsrude, C. D. Good, J. Ashburner, R. S. Frackowiak and C. D. Frith (2000). “Navigation-related structural change in the hippocampi of taxi drivers.” Proc Natl Acad Sci U S A 97(8): 4398–4403.

Mandai, K., T. Guo, C. St Hillaire, J. S. Meabon, K. C. Kanning, M. Bothwell and D. D. Ginty (2009). “LIG family receptor tyrosine kinase-associated proteins modulate growth factor signals during neural development.” Neuron 63(5): 614–627.

Martinowich, K., H. Manji and B. Lu (2007). “New insights into BDNF function in depression and anxiety.” Nat Neurosci 10(9): 1089–1093.

Mayseless, O., D. S. Berns, X. M. Yu, T. Riemensperger, A. Fiala and O. Schuldiner (2018). “Developmental Coordination during Olfactory Circuit Remodeling in Drosophila.” Neuron 99(6): 1204–1215 e1205.

McIlroy, G., I. Foldi, J. Aurikko, J. S. Wentzell, M. A. Lim, J. C. Fenton, N. J. Gay and A. Hidalgo (2013). “Toll-6 and Toll-7 function as neurotrophin receptors in the Drosophila melanogaster CNS.” Nat Neurosci 16(9): 1248–1256.

McLaughlin, C. N., I. V. Nechipurenko, N. Liu and H. T. Broihier (2016). “A Toll receptor-FoxO pathway represses Pavarotti/MKLP1 to promote microtubule dynamics in motoneurons.” J Cell Biol 214(4): 459–474.

McLaughlin, C. N., J. J. Perry-Richardson, J. C. Coutinho-Budd and H. T. Broihier (2019). “Dying Neurons Utilize Innate Immune Signaling to Prime Glia for Phagocytosis during Development.” Dev Cell 48(4): 506–522 e506.

Mehta, D., J. Voisey, D. Bruenig, W. Harvey, C. P. Morris, B. Lawford and R. M. Young (2018). “Transcriptome analysis reveals novel genes and immune networks dysregulated in veterans with PTSD.” Brain Behav Immun 74: 133–142.

Meyer, S. N., M. Amoyel, C. Bergantinos, C. de la Cova, C. Schertel, K. Basler and L. A. Johnston (2014). “An ancient defense system eliminates unfit cells from developing tissues during cell competition.” Science 346(6214): 1258236.

Minichiello, L. (2009). “TrkB signalling pathways in LTP and learning.” Nat Rev Neurosci 10(12): 850–860.

Nern, A., B. D. Pfeiffer and G. M. Rubin (2015). “Optimized tools for multicolor stochastic labeling reveal diverse stereotyped cell arrangements in the fly visual system.” Proc Natl Acad Sci U S A 112(22): E2967–2976.

Neve, K. A., J. K. Seamans and H. Trantham-Davidson (2004). “Dopamine receptor signaling.” J Recept Signal Transduct Res 24(3): 165–205.

Ohira, K. and M. Hayashi (2009). “A new aspect of the TrkB signaling pathway in neural plasticity.” Curr Neuropharmacol 7(4): 276–285.

Ohira, K., K. J. Homma, H. Hirai, S. Nakamura and M. Hayashi (2006). “TrkB-T1 regulates the RhoA signaling and actin cytoskeleton in glioma cells.” Biochem Biophys Res Commun 342(3): 867–874.

Okun, E., K. Griffioen, B. Barak, N. J. Roberts, K. Castro, M. A. Pita, A. Cheng, M. R. Mughal, R. Wan, U. Ashery and M. P. Mattson (2010). “Toll-like receptor 3 inhibits memory retention and constrains adult hippocampal neurogenesis.” Proc Natl Acad Sci U S A 107(35): 15625–15630.

Okun, E., K. J. Griffioen and M. P. Mattson (2011). “Toll-like receptor signaling in neural plasticity and disease.” Trends Neurosci 34(5): 269–281.

Pare, A. C., A. Vichas, C. T. Fincher, Z. Mirman, D. L. Farrell, A. Mainieri and J. A. Zallen (2014). “A positional Toll receptor code directs convergent extension in Drosophila.” Nature 515(7528): 523–527.

Park, H. and M. M. Poo (2013). “Neurotrophin regulation of neural circuit development and function.” Nat Rev Neurosci 14(1): 7–23.

Patel, V., A. M. Patel and J. J. McArdle (2016). “Synaptic abnormalities of mice lacking toll-like receptor (TLR)-9.” Neuroscience 324: 1–10.

Pech, U., N. H. Revelo, K. J. Seitz, S. O. Rizzoli and A. Fiala (2015). “Optical dissection of experience-dependent pre- and postsynaptic plasticity in the Drosophila brain.” Cell Rep 10(12): 2083–2095.

Placais, P. Y., E. de Tredern, L. Scheunemann, S. Trannoy, V. Goguel, K. A. Han, G. Isabel and T. Preat (2017). “Upregulated energy metabolism in the Drosophila mushroom body is the trigger for long-term memory.” Nat Commun 8: 15510.

Placais, P. Y., S. Trannoy, G. Isabel, Y. Aso, I. Siwanowicz, G. Belliart-Guerin, P. Vernier, S. Birman, H. Tanimoto and T. Preat (2012). “Slow oscillations in two pairs of dopaminergic neurons gate long-term memory formation in Drosophila.” Nat Neurosci 15(4): 592–599.

Poo, M. M. (2001). “Neurotrophins as synaptic modulators.” Nat Rev Neurosci 2(1): 24–32.

Riemensperger, T., G. Isabel, H. Coulom, K. Neuser, L. Seugnet, K. Kume, M. Iche-Torres, M. Cassar, R. Strauss, T. Preat, J. Hirsh and S. Birman (2011). “Behavioral consequences of dopamine deficiency in the Drosophila central nervous system.” Proc Natl Acad Sci U S A 108(2): 834–839.

Riemensperger, T., A. R. Issa, U. Pech, H. Coulom, M. V. Nguyen, M. Cassar, M. Jacquet, A. Fiala and S. Birman (2013). “A single dopamine pathway underlies progressive locomotor deficits in a Drosophila model of Parkinson disease.” Cell Rep 5(4): 952–960.

Rolls, A., R. Shechter, A. London, Y. Ziv, A. Ronen, R. Levy and M. Schwartz (2007). “Toll-like receptors modulate adult hippocampal neurogenesis.” Nat Cell Biol 9(9): 1081–1088.

Sachse, S., E. Rueckert, A. Keller, R. Okada, N. K. Tanaka, K. Ito and L. B. Vosshall (2007). “Activity-dependent plasticity in an olfactory circuit.” Neuron 56(5): 838–850.

Sahu, M. P., Y. Pazos-Boubeta, C. Pajanoja, S. Rozov, P. Panula and E. Castren (2019). “Neurotrophin receptor Ntrk2b function in the maintenance of dopamine and serotonin neurons in zebrafish.” Sci Rep 9(1): 2036.

Shafer, O. T., D. J. Kim, R. Dunbar-Yaffe, V. O. Nikolaev, M. J. Lohse and P. H. Taghert (2008). “Widespread receptivity to neuropeptide PDF throughout the neuronal circadian clock network of Drosophila revealed by real-time cyclic AMP imaging.” Neuron 58(2): 223–237.

Sharma, A. (2012). “Genome-wide expression analysis in epilepsy: a synthetic review.” Curr Top Med Chem 12(9): 1008–1032.

Squillace, S. and D. Salvemini (2022). “Toll-like receptor-mediated neuroinflammation: relevance for cognitive dysfunctions.” Trends Pharmacol Sci 43(9): 726–739.

Stoilov, P., E. Castren and S. Stamm (2002). “Analysis of the human TrkB gene genomic organization reveals novel TrkB isoforms, unusual gene length, and splicing mechanism.” Biochem Biophys Res Commun 290(3): 1054–1065.

Sugie, A., S. Hakeda-Suzuki, E. Suzuki, M. Silies, M. Shimozono, C. Mohl, T. Suzuki and G. Tavosanis (2015). “Molecular Remodeling of the Presynaptic Active Zone of Drosophila Photoreceptors via Activity-Dependent Feedback.” Neuron 86(3): 711–725.

Sun, J., A. Q. Xu, J. Giraud, H. Poppinga, T. Riemensperger, A. Fiala and S. Birman (2018). “Neural Control of Startle-Induced Locomotion by the Mushroom Bodies and Associated Neurons in Drosophila.” Front Syst Neurosci 12: 6.

Sur, M. and J. L. Rubenstein (2005). “Patterning and plasticity of the cerebral cortex.” Science 310(5749): 805–810.

Sutcliffe, B., M. G. Forero, B. Zhu, I. M. Robinson and A. Hidalgo (2013). “Neuron-type specific functions of DNT1, DNT2 and Spz at the Drosophila neuromuscular junction.” PLoS One 8(10): e75902.

Talay, M., E. B. Richman, N. J. Snell, G. G. Hartmann, J. D. Fisher, A. Sorkac, J. F. Santoyo, C. Chou-Freed, N. Nair, M. Johnson, J. R. Szymanski and G. Barnea (2017). “Transsynaptic Mapping of Second-Order Taste Neurons in Flies by trans-Tango.” Neuron 96(4): 783–795 e784.

Tamada, M., J. Shi, K. S. Bourdot, S. Supriyatno, K. H. Palmquist, O. L. Gutierrez-Ruiz and J. A. Zallen (2021). “Toll receptors remodel epithelia by directing planar-polarized Src and PI3K activity.” Dev Cell 56(11): 1589–1602 e1589.

Tauszig, S., E. Jouanguy, J. A. Hoffmann and J. L. Imler (2000). “Toll-related receptors and the control of antimicrobial peptide expression in Drosophila.” Proc Natl Acad Sci U S A 97(19): 10520–10525.

Technau, G. M. (1984). “Fiber number in the mushroom bodies of adult Drosophila melanogaster depends on age, sex and experience.” J Neurogenet 1(2): 113–126.

Tempel, B. L., N. Bonini, D. R. Dawson and W. G. Quinn (1983). “Reward learning in normal and mutant Drosophila.” Proc Natl Acad Sci U S A 80(5): 1482–1486.

Tessarollo, L. and S. Yanpallewar (2022). “TrkB Truncated Isoform Receptors as Transducers and Determinants of BDNF Functions.” Front Neurosci 16: 847572.

Ulian-Benitez, S., S. Bishop, I. Foldi, J. Wentzell, C. Okenwa, M. G. Forero, B. Zhu, M. Moreira, M. Phizacklea, G. McIlroy, G. Li, N. J. Gay and A. Hidalgo (2017). “Kek-6: A truncated-Trk-like receptor for Drosophila neurotrophin 2 regulates structural synaptic plasticity.” PLoS Genet 13(8): e1006968.

Vahid-Ansari, F. and P. R. Albert (2021). “Rewiring of the Serotonin System in Major Depression.” Front Psychiatry 12: 802581.

Vaughen, J. P., E. Theisen, I. M. Rivas-Serna, A. B. Berger, P. Kalakuntla, I. Anreiter, V. C. Mazurak, T. P. Rodriguez, J. D. Mast, T. Hartl, E. O. Perlstein, R. J. Reimer, M. T. Clandinin and T. R. Clandinin (2022). “Glial control of sphingolipid levels sculpts diurnal remodeling in a circadian circuit.” Neuron 110(19): 3186–3205 e3187.

Waddell, S. (2013). “Reinforcement signalling in Drosophila; dopamine does it all after all.” Curr Opin Neurobiol 23(3): 324–329.

Wang, C. S., E. T. Kavalali and L. M. Monteggia (2022). “BDNF signaling in context: From synaptic regulation to psychiatric disorders.” Cell 185(1): 62–76.

Ward, A., W. Hong, V. Favaloro and L. Luo (2015). “Toll receptors instruct axon and dendrite targeting and participate in synaptic partner matching in a Drosophila olfactory circuit.” Neuron 85(5): 1013–1028.

Weber, A. N., S. Tauszig-Delamasure, J. A. Hoffmann, E. Lelievre, H. Gascan, K. P. Ray, M. A. Morse, J. L. Imler and N. J. Gay (2003). “Binding of the Drosophila cytokine Spatzle to Toll is direct and establishes signaling.” Nat Immunol 4(8): 794–800.

Wiesel, T. N. (1982). “Postnatal development of the visual cortex and the influence of environment.” Nature 299(5884): 583–591.

Wohleb, E. S., T. Franklin, M. Iwata and R. S. Duman (2016). “Integrating neuroimmune systems in the neurobiology of depression.” Nat Rev Neurosci 17(8): 497–511.

Woollett, K. and E. A. Maguire (2011). “Acquiring “the Knowledge” of London’s layout drives structural brain changes.“ Curr Biol 21(24): 2109–2114.

Yacoubian, T. A. and D. C. Lo (2000). “Truncated and full-length TrkB receptors regulate distinct modes of dendritic growth.” Nat Neurosci 3(4): 342–349.

Yang, T., Z. Nie, H. Shu, Y. Kuang, X. Chen, J. Cheng, S. Yu and H. Liu (2020). “The Role of BDNF on Neural Plasticity in Depression.” Front Cell Neurosci 14: 82.

Zhang, J., T. Tang, R. Zhang, L. Wen, X. Deng, X. Xu, W. Yang, F. Jin, Y. Cao, Y. Lu and X. Q. Yu (2024). “Maintaining Toll signaling in Drosophila brain is required to sustain autophagy for dopamine neuron survival.” iScience 27(2): 108795.

Zhu, B., J. A. Pennack, P. McQuilton, M. G. Forero, K. Mizuguchi, B. Sutcliffe, C. J. Gu, J. C. Fenton and A. Hidalgo (2008). “Drosophila neurotrophins reveal a common mechanism for nervous system formation.” PLoS Biol 6(11): e284.

