## Supporting Information for "A neurotrophin functioning with a Toll regulates structural plasticity in a dopaminergic circuit"

SUPPLEMENTARY FIGURE S1 Identification of neurotransmitter type for DNT- 2A neurons.

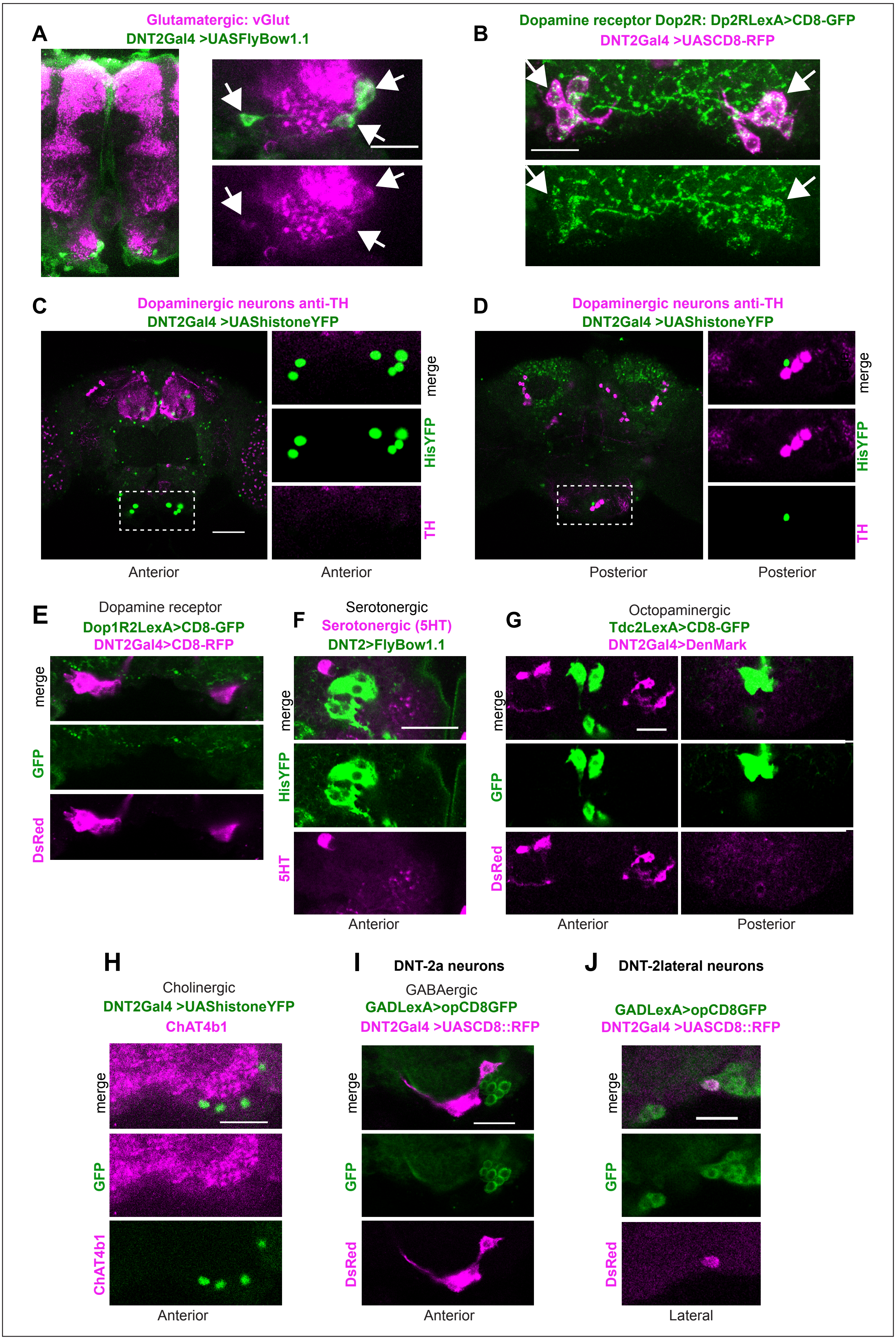

Supplementary Figure S2 Identification of Toll-6 and kek-6 expressing neurons

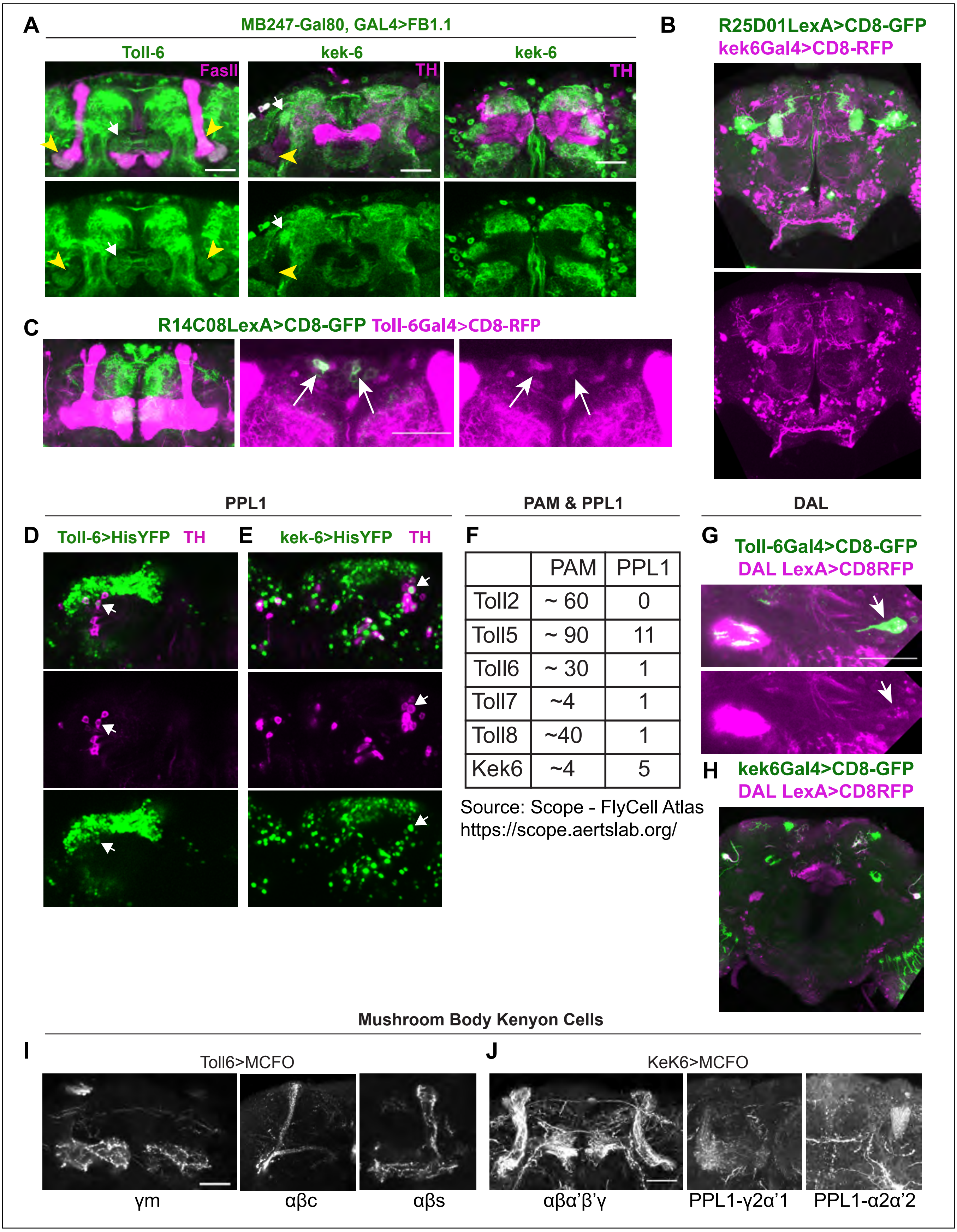


| | PAM | PPL1 |
|---|---|---|
| Toll2 | ~ 60 | 0 |
| Toll5 | ~ 90 | 11 |
| Toll6 | ~ 30 | 1 |
| Toll7 | ~4 | 1 |
| Toll8 | ~40 | 1 |
| Kek6 | ~4 | 5 |

Source: Scope - FlyCell Atlas
https://scope.aertslab.org/

Figure S3   TransTango and connectivity controls.

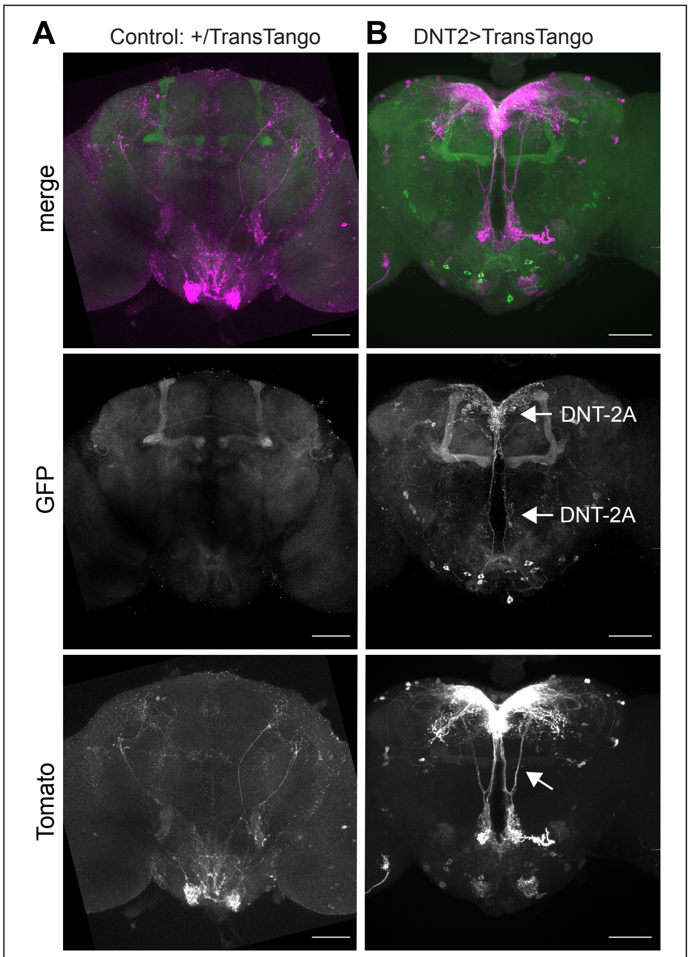


controls for Figure 2A

Supplementary Figure S4
Neuronal activity increased production and cleavage of DNT-2

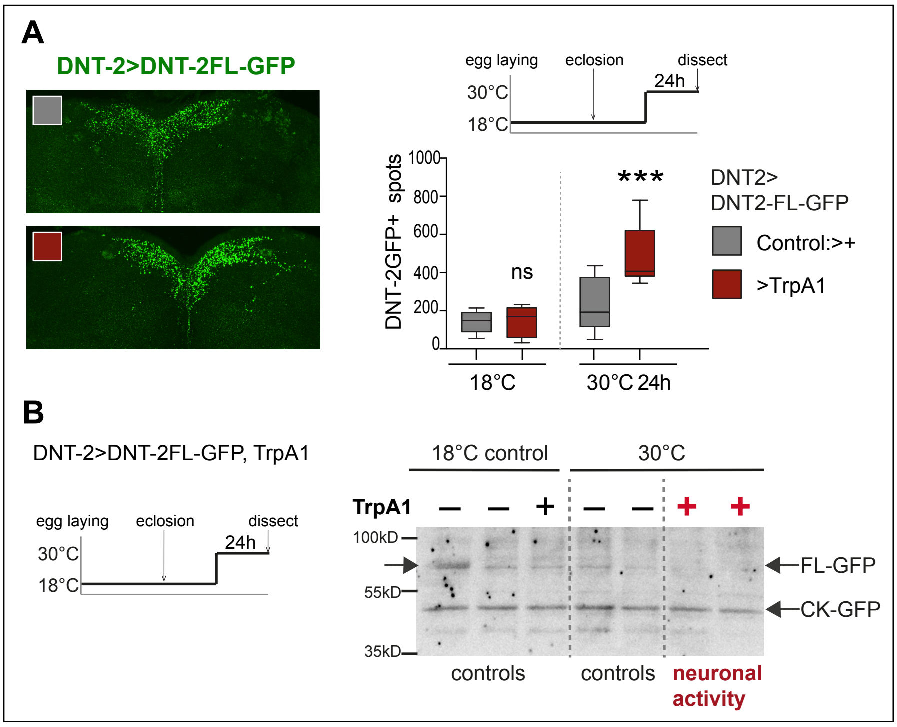

Figure S5        Controls for Figure 5B

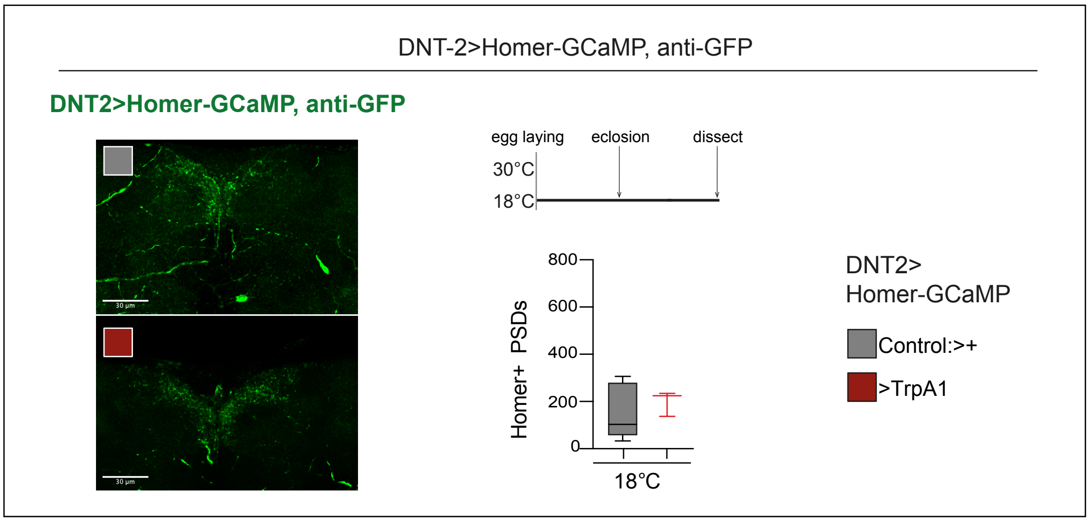

Figure S6 Locomotion Controls

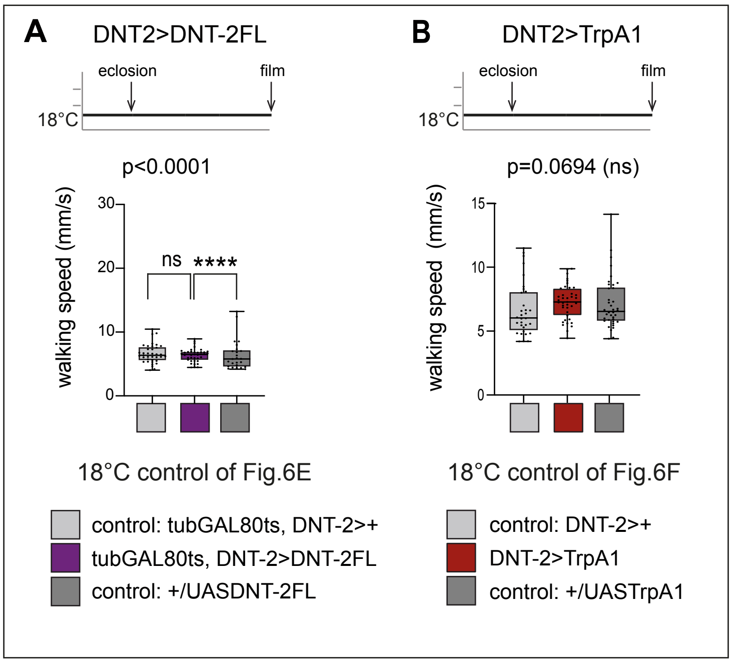

Supplementary Figure S7
Alterating *DNT-2* levels induces seizures

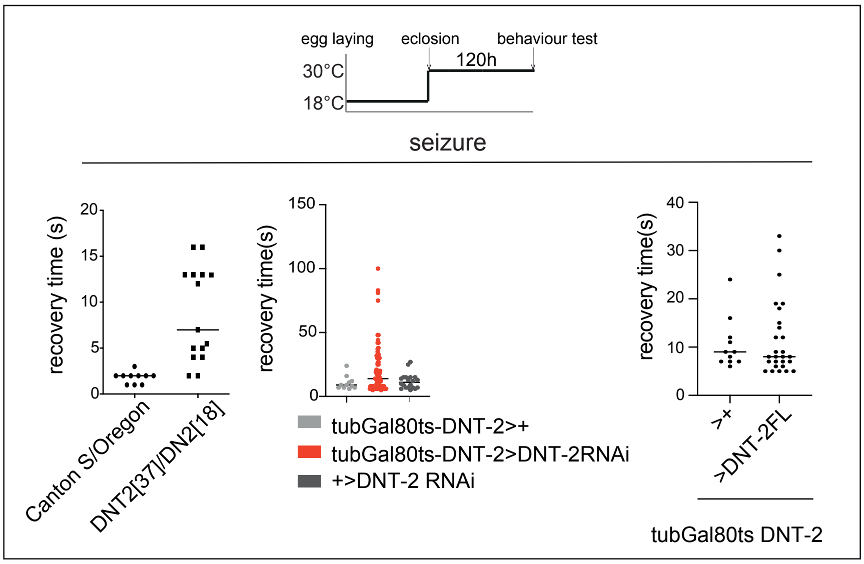
